# Single-cell analysis of basal cell carcinoma reveals heat shock proteins promote tumor growth in response to WNT5A-mediated inflammatory signals

**DOI:** 10.1101/2021.10.07.463571

**Authors:** Christian F. Guerrero-Juarez, Gun Ho Lee, Yingzi Liu, Shuxiong Wang, Yutong Sha, Rachel Y. Chow, Tuyen T.L. Nguyen, Sumaira Aasi, Matthew Karikomi, Michael L. Drummond, Qing Nie, Kavita Sarin, Scott X. Atwood

**Affiliations:** Department of Developmental and Cell Biology, University of California, Irvine, Irvine, CA, 92697, USA; Department of Mathematics, University of California, Irvine, Irvine, CA 92697, USA; NSF-Simons Center for Multiscale Cell Fate Research, University of California, Irvine, Irvine, CA 92697, USA; Center for Complex Biological Systems, University of California, Irvine, Irvine, CA 92697, USA; Department of Dermatology, Stanford University School of Medicine, Stanford, CA 94305, USA; Department of Dermatology, University of California, Irvine, Irvine, CA, 92697, USA; Chao Family Comprehensive Cancer Center, University of California, Irvine, Irvine, CA, 92697, USA

## Abstract

How basal cell carcinoma (BCC) interacts with its tumor microenvironment to promote growth is unclear. Here we use singe-cell RNA sequencing to define the human BCC ecosystem and discriminate between normal and malignant epithelial cells. We identify spatial biomarkers of both tumors and their surrounding stroma that reinforce the heterogeneity of each tissue type. Combining pseudotime, RNA velocity, cellular entropy, and regulon analysis in stromal cells reveal a cancer-specific rewiring of fibroblasts where STAT1, TGF-β, and inflammatory signals induce a non-canonical WNT5A program that maintains the stromal inflammatory state. Cell-cell communication modeling suggests that tumors respond to the sudden burst of fibroblast-specific inflammatory signaling pathways by producing heat shock proteins, which we validated *in situ*. Finally, dose-dependent treatment with an HSP70 inhibitor suppresses *in vitro* BCC cell growth and Hedgehog signaling and *in vivo* tumor growth in a BCC mouse model, validating HSP70’s essential role in tumor growth and reinforcing the critical nature of tumor microenvironment crosstalk in BCC progression.

## Introduction

Basal cell carcinoma (BCC) is a locally invasive skin cancer and the most common human cancer worldwide (Cameron et al., 2019a) with an estimated lifetime risk between 20-30% (Rigel et al., 1996) and increasing incidence rates in a number of regions including North America (Mudigonda et al., 2013; Nugent et al., 2005), Europe (de Vries et al., 2004; Rudolph et al., 2015), Asia (Koh et al., 2003; Sng et al., 2009), and Australia (Staples et al., 2006). BCCs originate from inappropriate activation of the Hedgehog (HH) signaling pathway, in which secreted HH ligand binds the cholesterol transporter Patched Homologue 1 (PTCH1) and negates PTCH1-mediated suppression of the G-protein coupled receptor Smoothened (SMO). SMO then activates the GLI family of transcription factors to promote proliferation and tumor growth (Bakshi et al., 2017; Cameron et al., 2019a; Rubin et al., 2005).Although the mortality rate for BCC is low (Wehner et al., 2018), the large affected patient population imposes significant morbidity and cost (Rees et al., 2015; Rowell et al., 2016; Wysong et al., 2013).

Although surgery remains the gold standard of therapy for BCC (Cameron et al., 2019b; Clark et al., 2014), it is not a practical option for tumors on cosmetically sensitive body parts or for metastatic disease. SMO inhibitors, vismodegib and sonidegib, have emerged as a promising treatments for advanced disease (Basset-Seguin et al., 2015; Tang et al., 2019), with a response rate around 30% in metastatic BCC and 45% in locally advanced BCC (Chang et al., 2014; Sekulic et al., 2013; Sekulic et al., 2012). However, SMO mutations driving drug resistance is common (Atwood et al., 2015; Sharpe et al., 2015) and up to 21% of patients treated with vismodegib were found to undergo tumor regrowth during treatment (Chang and Oro, 2012). Additional pathways that contribute to BCC drug resistance include PI3K/MTOR (Chow et al., 2021a; Chow et al., 2021b), WNT (Sanchez-Danes et al., 2018), aPKC iota/lambda (Atwood et al., 2013), NOTCH1 (Eberl et al., 2018), RAS/MAPK (Kuonen et al., 2019; Zhao et al., 2015), and activation of MRTF (Whitson et al., 2018; Yao et al., 2020), to name a few. New therapeutic options are needed to treat advanced BCC.

How stroma interact with, and promote the growth of, BCCs is unclear. Upon hierarchical clustering of cancer-associated fibroblast (CAF) markers in BCC, squamous cell carcinoma, and melanoma, three distinct subgroups can be stratified, each corresponding to the specific cancer type (Sasaki et al., 2018). Specifically, BCC CAFs are notable for their high expression of PDGFR-beta, S100A4, and TWIST. Within different histopathologic subtypes of BCCs, the tumor-to-stroma ratio is significantly divergent, with infiltrative BCCs presenting the lowest ratio (Lesack and Naugler, 2012). Genes coding for extracellular matrix components are also upregulated in BCCs, suggesting a tumor-induced remodeling of the stromal matrix (Omland et al., 2017). Additionally, expression of stromal proteins have been shown to predict the aggressiveness of BCCs (Adegboyega et al., 2010) and distinguish between infiltrative BCC and desmoplastic trichoepithelioma (Abbas et al., 2010). Together, these studies show expressed factors in BCC-stroma can play important roles in tumor growth, angiogenesis, and metastasis (Bhowmick et al., 2004; Lynch and Matrisian, 2002). Defining BCC-stroma interactions may be a vital, yet understudied, part of tumor progression and result in more efficacious therapies.

Single-cell RNA sequencing (scRNA-seq) technologies allow for the analysis of intra-sample heterogeneity, tumor/sample microenvironment, pathogenic pathways, and cell-cell interactions in oncogenic contexts (Gonzalez-Silva et al., 2020). Using this technology, we define BCC cellular heterogeneity, cell-cell interactions, and novel active pathways in BCC. We differentiate between malignant and normal epithelia, identify a stromal inflammatory response driven by WNT5A, characterize a subgroup of BCC keratinocytes that overexpress heat shock proteins, and provide data supporting the HSP pathway as a potential novel therapeutic target for BCC.

## Results

### Cellular ecosystem of human BCC

To define the cellular ecosystem of human BCCs, we sorted viable, single cells *in toto* from primary human BCC surgical discards (n = 4), including peri-tumor normal skin (PTS) tissues (n = 2), and subjected them to 3’-droplet-enabled scRNA-seq (**Fig. 1A; Supplementary Fig. 1**) (Macosko et al., 2015). The primary BCC subtypes considered in this study included superficial, nodular and infiltrative BCC (ID: BCC-I; k = 9,837 cells); superficial and nodular BCC (ID: BCC-II; k = 11,724 cells); unknown/”hybrid” BCC (ID: BCC-III; k = 6,712 cells); and infiltrative with perineural invasion BCC (ID: BCC-IV; k = 8,569 cells). PTS tissues constituted skin directly adjacent to BCC lesions. In total, we processed 56,162 raw single cells (k_PTS_ = 17,*727* versus k_BCC_ = 38,435). Following putative doublet/multiplet removal and quality control filtering of individual libraries (**Supplementary Fig. 2; Supplementary Table 1-3**) (Fan et al., 2016; Satija et al., 2015; Wolock et al., 2019), 52,966 ‘valid’ cells remained (k_PTS_ = 16,903 versus k_BCC_ = 37,667). To resolve the cellular diversity present in individual tumors and enable downstream query and comparative gene expression analyses, we processed and characterized individual BCCs and using Seurat (Stuart et al., 2019) and visualized the inferred putative cell types. We identified ten, coarse-grained cell types which included *MKI67*^+^ proliferative epithelial cells, *KRT14*^+^ basal epithelial/tumor cells, terminally differentiated *IVL*^+^ keratinocytes, *AZPG*^+^ appendage-associated cells, *PDGFRA*^+^ fibroblastic cells, *RGS5*^+^ fibroblast-like cells, *TIE1*^+^ endothelial cells, *PROX1*^+^ lymphatic endothelial cells, *MLANA*^+^ melanocytic cells, and immune cells identified by expression of *PTPRC* (**Fig. 1B**). We did not confidently identify cell clusters with gene expression signatures enriched for *Stratum spinosum* keratinocytes or Schwann/neural-like cells (**Supplementary Fig. 3**).

**Figure 1.**
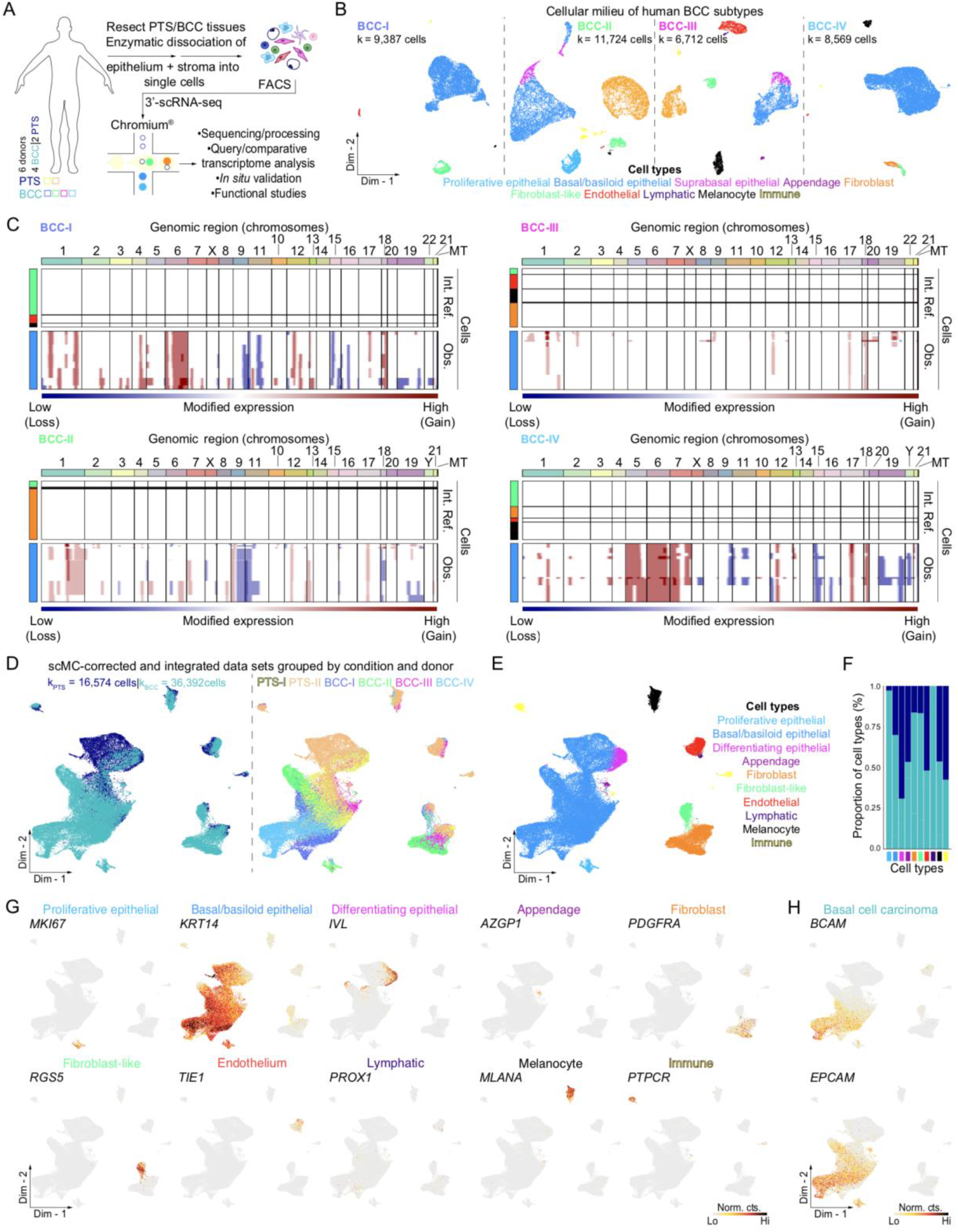
Cellular characterization of human basal cell carcinoma subtypes using single-cell RNA-sequencing. **A**. Schematic representation of *in toto* epithelial and stromal tissue isolation and processing from human peri-tumor skin (PTS) and basal cell carcinoma (BCC) tissues for 3’-droplet-enabled single-cell RNA-sequencing (scRNA-seq). **B**. Two-dimensional clustering of single cells isolated from individual human basal cell carcinoma subtypes. IDs represent subtype and donor. BCC subtypes are color-coded based on subtype and donor and include: superficial, nodular, and infiltrative (BCC-I); superficial and nodular (BCC-II); unknown/”hybrid” (BCC-III); and infiltrative with perineural invasion (BCC-IV). Ten distinct meta-clusters are identified at distinct proportions across BCC subtypes and annotated with their putative cell type identities. The putative identity of each cell meta-cluster is defined on the bottom and color-coded accordingly. **C**. Copy number variant analysis of putative malignant epithelial cells with InferCNV. Dark blue indicates low modified expression – corresponding to genomic loss; dark red indicates high modified gene expression – corresponding to genomic gain. Internal reference cells refer to non-epithelial, non-immune cells. Internal reference refers to non-epithelial control cells. Observations refer to putative malignant epithelial cells. Genomic regions (chromosomes) are labeled and color-coded. **D**. Clustering of corrected and integrated PTS and BCC data sets grouped by condition and donor using scMC. Conditions and donor are labeled and color-coded. **E**. Two-dimensional clustering reveals cellular heterogeneity of integrated human PTS and BCC data sets. Ten distinct meta-clusters are identified at various proportions across BCC subtypes and annotated with their putative cell type identities. The putative identity of each cell meta-cluster is defined on the right and color-coded accordingly per cell type. **F**. Proportion of cell types grouped by condition. **G, H**. Feature plots showing *bona fide* genes (**G**) and BCC-specific epithelial markers (**H**). Light gray indicates low normalized gene expression based on normalized counts; black indicates high normalized gene expression based on normalized counts. Abbreviations: Int. Ref. – internal reference; Obs. – observations; MT – mitochondrial; FACS – fluorescent activated cell sorting; Norm. cts. – normalized counts.

To identify putative malignant tumor cells present in primary BCC samples, we subjected the *KRT14*^+^ epithelial/tumor cells to InferCNV analysis (Patel et al., 2014; Puram et al., 2017; Tirosh et al., 2016a; Tirosh et al., 2016b). We observed aberrant genomic profiles, associated with chromosome duplication (dark red) and deletion (dark blue), in *KRT14*^+^ epithelial cells from BCC-I, BCC-II, and BCC-IV donors when compared to their counterpart non-epithelial, non-immune internal reference cells (**Fig. 1C**). Interestingly, BCC-III did not display significant aberrant genomic structure changes when compared to other BCC subtypes. Rather, its profile resembled more those from non-appendage, *KRT14*^+^ epithelial cells present in the PTS samples (**Supplementary Fig. 4**), suggesting that some tumors do not have significant copy number variations driving tumor growth. Although InferCNV inferred aberrant genomic changes in *KRT14*^+^ epithelial cells, it cannot identify individual malignant cells. We were also not able to confidently identify individual malignant cells using copyKAT (Gao et al., 2021), even when using non-epithelial cells as reference (data not shown).

When integrating both BCC and PTS datasets using Seurat, we noticed independent clustering of BCC *KRT14*^+^ epithelial/tumor cells from PTS, with further inter-BCC partitioning (**Supplementary Fig. S5A**). Unlike *KRT14*^+^ epithelial/tumor cells, all other non-epithelial cell types did not drift or cluster independently from each other regardless of donor. The high BCC tumor heterogeneity is in congruence with other reports indicating a high degree of transcriptome-driven epithelial, inter-tumoral heterogeneity in other human cancers, including melanoma and squamous cell carcinoma (Ji et al., 2020; Nguyen et al., 2018; Puram et al., 2017; Tirosh et al., 2016a; Yost et al., 2019). To determine the best approach to identify BCC-associated *KRT14*^+^ epithelial/tumor cell states that significantly differ from PTS, we compared Seurat-based integration with four distinct yet widely popular methods, including SCTransform (Hafemeister and Satija, 2019; Satija et al., 2015; Stuart et al., 2019), LIGER (Liu et al., 2020; Welch et al., 2019), Harmony (Korsunsky et al., 2019), and scMC (Zhang and Nie, 2021) (**Fig. 1D; Supplementary Fig. 5**). All algorithms clustered non-epithelial cells together, irrespective of condition or donor. However, Seurat, SCTransform, LIGER, and Harmony clustered epithelial cells indistinctly, irrespective of condition or donor; whereas clustering with Seurat was driven entirely by donor, making it difficult to identify and interpret BCC-specific epithelial cell types or states (**Supplementary Fig. 5A-D**). In sharp contrast, scMC clustered BCC and PTS epithelial cells distinctly, while maintaining clustering of transcriptionally similar cell types (**Fig. 1D**). As scMC retains biological variation while removing technical variation associated with each sample, we therefore used the resultant scMC-corrected BCC-PTS data for downstream query and comparative analysis.

scMC maintained the same ten distinct cell types found by independent BCC clustering with Seurat (**Fig. 1E-G**). Quantification of each coarse-grained cell type partitioned by condition and donor revealed relative cell type frequency similarities across BCC and PTS (**Fig. 1F**). Non-PTS *KRT14*^+^ epithelial/tumor cells uniquely expressed known BCC-associated gene biomarkers including *BCAM* and *EPCAM* (**Fig. 1H**) (Yost et al., 2019). Interestingly, *KRT14*^+^ epithelial/tumor cells from BCC-III showed a “hybrid” position where many cells significantly overlapped with the PTS samples, whereas other cells uniquely clustered singly, matching our previous observations with InferCNV analyses (**Fig. 1C-D**). In sum, our benchmarking approach and comparative scRNA-seq clustering analyses resolved the distinct cellular landscape of human BCCs and revealed major *KRT14*^+^ epithelial cell type differences compared to PTS, suggesting a high level of inter- and intra-tumor transcriptional heterogeneity between human BCC samples.

### Defining normal versus malignant epithelial cells

To define the epithelial/tumor cellular landscape of human BCC and PTS samples, we sub-clustered 30,058 *KRT14*^+^ epithelial-derived cells (k_PTS_ = 5,146 versus k_BCC_ = 24,872) and identified fifteen coarse-grained epithelial cell clusters, all defined by expression of unique gene biomarkers (**Fig. 2A-C**). Three of the subpopulations (IFE I-III) appear to be normal epithelia and make up nearly all the PTS samples and a small proportion of the BCC samples, whereas the rest of the cells cluster uniquely to the BCC-associated samples (BAS I-XII). Our integrative comparative analysis did not reveal the presence of unique Tumor Specific Keratinocytes (TSKs), as has been previously reported for squamous cell carcinoma (Ji et al., 2020), but rather BCC-associated keratinocytes that have common and unique gene expression patterns. To explore their gene expression profile and spatial architecture, we spatially resolved select gene-products, including *KRT15* (BAS-II), *LHX2* (BAS-IV), and *ACTA2* (BAS-XI). KRT15 marked a subset of KRT14^high^ tumor nests (**Fig. 2D**), LHX2 marked the nucleus of cells along the outer periphery of KRT14^high^ tumor nests (**Fig. 2E**), and ACTA2 marked the outer periphery of KRT14^low^ tumor nests (**Fig. 2F**), reinforcing the heterogeneity of BCC tumors.

**Figure 2.**
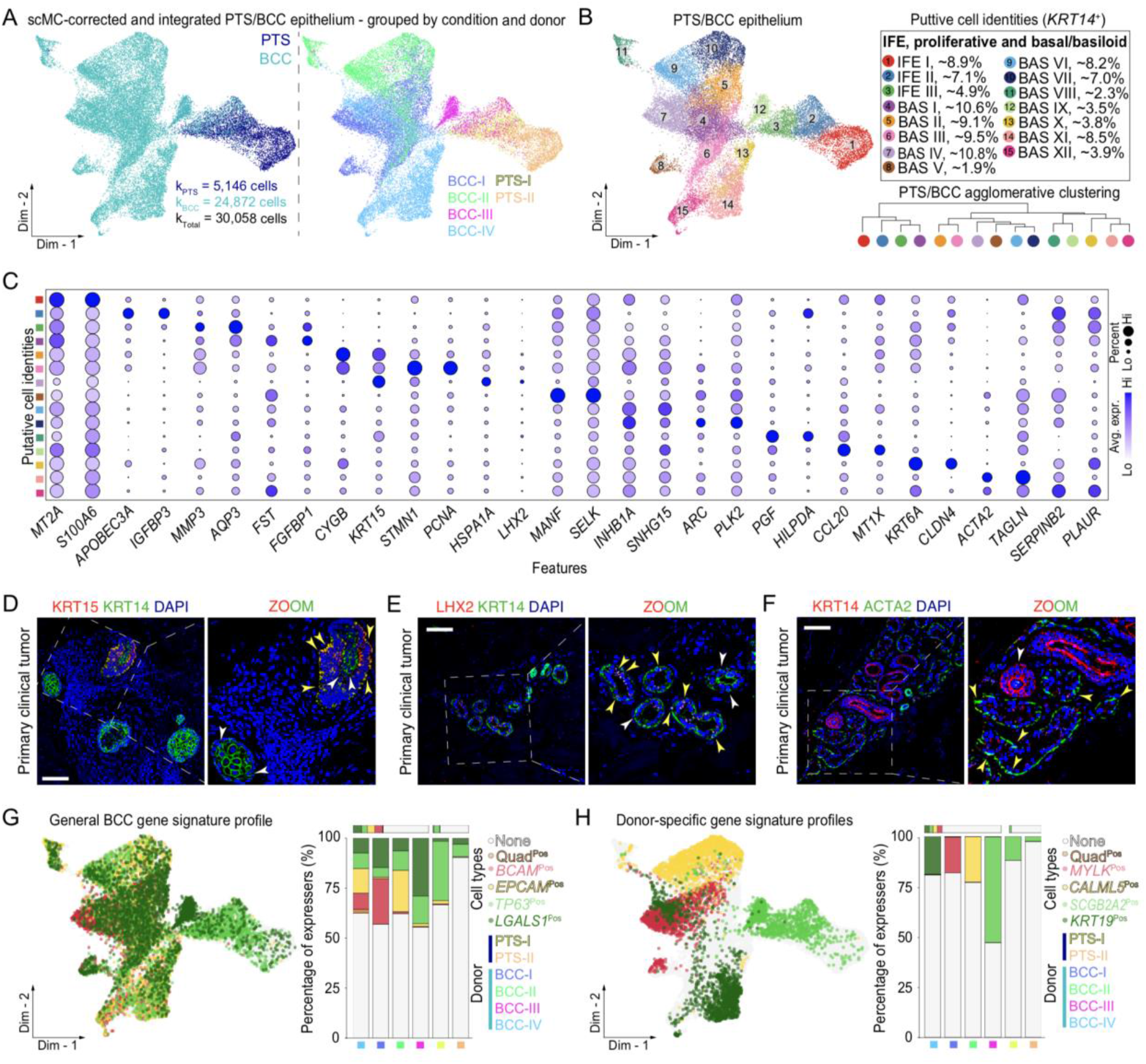
Comparison of epithelial cells reveals regulators of malignancy. **A-B**. Clustering of 30,058 batch-corrected *KRT14*^+^ epithelial cells from peri-tumor skin (PTS) and basal cell carcinoma (BCC) subtypes grouped by condition and donor. Fifteen putative *KRT14*^+^ epithelial cell identities, including one proliferating epithelial and three interfollicular epithelial cells, and eleven basal/basaloid epithelial cells were identified, and their identity defined on the right. PTS/BCC agglomerative clustering shows relationships between *KRT14*^+^ epithelial cells. Cells are color-coded accordingly. **C**. Dotplot of top two marker genes identified by differential gene expression amongst epithelial cells. Light gray indicates low average gene expression; purple indicates high average gene expression. Size of circle represents the percentage of cells expressing gene markers of interest. **D-F**. Immunostaining of select BCC-epithelial cell markers show cluster-specificity and distinct spatial localization in primary clinical tumors. Sections were counterstained with DAPI. Inset shows magnified area in BCC nest. White arrows point at epithelial cells expressing gene of interest. Yellow arrows point at epithelial cells expressing gene of interest and KRT14 Size bars: 100 µm. **G**. General BCC signature profile overlaid on two-dimensional epithelial cell embedding. Percentage of cells expressing *BCAM, EPCAM, TP63*, and *LGALS1*. Quadrupled positive cells are labeled orange and negative cells are labeled light gray. **H**. Donor-specific BCC gene signature profile overlaid on two-dimensional epithelial cell embedding. Percentage of cells expressing *MYLK, CALML5, SCGB2A2*, and *KRT19*. Quadrupled positive cells are labeled orange and negative cells are labeled light gray. Differentially expressed genes per BCC subtype considered unique and significant if 50% of cells in each condition express the gene at a Log_2_ fold change of 0.25X (Wilcoxon Rank Sum Test, P_adj_ < 0.05).

We next defined the identity of ‘transformed’ cells by scoring individual *KRT14*^+^ cells with a previously defined BCC-associated gene expression profile that includes co-expression of *EPCAM, BCAM*, and *TP63* (Bernemann et al., 2000; Bircan et al., 2006; Depianto et al., 2010; Tellechea et al., 1993). When grouped by condition and quantified, we observed that the majority of BCC-associated cells expressed some of these markers but not all three. Of interest, *EPCAM* and *BCAM* were rather unique to BCC, whereas *TP63* was lowly expressed in PTS samples. To develop a better measure of transformation, we identified prominent gene expression differences between BCC and PTS epithelial cells. This approach led to the identification of *LGALS1* as a gene that is highly upregulated in BCC epithelial cells (**Fig. 2G**). LGALS1 has been previously implicated in pancreatic ductal adenocarcinoma (Orozco et al., 2018), clear cell renal cell carcinoma (White et al., 2014), cervical cancer (Chetry et al., 2020), and malignant melanomas (Bolander et al., 2008). We conducted a similar approach to identify genes associated with the different BCC samples in our cohort of donors. We identified *MYLK, CALM5, SCGB2A2*, and *KRT19* as highly expressed within each tumor sample (**Fig. 2H**).

We were interested in using scRNA-seq data to determine whether we could match genes or gene-specific loci identified in other bulk transcriptome of genomic studies with individual cells within the BCC macro-environment. To do this, we used our human BCC scRNA-seq data and overlaid expression of BCC risk-loci identified via GWAS studies in BCC (Chahal et al., 2016) (**Supplementary Fig. 6**). We successfully identified several differentially expressed genes that were associated with specific BCC risk-loci and that were expressed only in BCC epithelial cells, including *BNC2, CUX1, ZBTB10*, and *CASC15* (**Supplementary Fig. 6**). We also employed a similar approach to identify vismodegib-resistant genes from bulk-level RNA-seq analysis of advanced BCC (Atwood et al., 2015) that are expressed only in BCC epithelial cells (**Supplementary Fig. 7A-B**). We found a cohort of genes that were non-specific and had broad expression in other cell types, including *SLC39A14, DUSP10*, and *SOX4* (**Supplementary Fig. 7C**). In contrast, other genes displayed unique expression in BCC epithelial cells, including *FBN3, SH3GL3*, and *KC6* (**Supplementary Fig. 7C**). This approach could enable the identification of genes specific to certain BCC subtypes, or BCC epithelial subclusters.

### RNA dynamics show de-differentiation in BCC

As BCCs display both inter- and intra-tumor heterogeneity, we performed RNA velocity analysis using single cell Velocity (scVelo) (Bergen et al., 2020a; La Manno et al., 2018) to better estimate and generalize transient cell states within *KRT14*^+^ epithelial cells. Coupling RNA velocity vectors with Markovnikov root and terminal states suggested that superficial and nodular (BCC-I); superficial, nodular, and infiltrative (BCC-II); and the unknown/”hybrid” subtype BCC (BCC-III) demonstrated velocity vectors pointing toward a terminal state associated with high levels of BCC-associated signature genes and high in HH and WNT pathway genes (**Supplementary Fig. 8A-C**). In contrast, the infiltrative with perineural invasion BCC (BCC-IV) displayed vectors pointing away from a region high in HH genes (**Supplementary Fig. 8D**). Interestingly, velocities derived from cells with a clear high late differentiation gene signature (Lopez-Pajares et al., 2015) in BCC-I and BCC-II suggests a potential de-differentiation fate choice of late differentiation epithelial cells in favor of a more basal-like fate in BCC (**Supplementary Fig. 8A-B**), in contrast to normal epithelia which display velocities going toward the high late differentiation gene signature (**Supplementary Fig. 8C**).

### Fibroblast heterogeneity and function in human BCC

Recent studies have identified a large degree of functional heterogeneity in fibroblasts (FIB) and fibroblast-like (FIB-like) cells across different states and conditions in human (Ji et al., 2020; Philippeos et al., 2018; Puram et al., 2017; Sole-Boldo et al., 2020; Stephenson et al., 2018; Tirosh et al., 2016a) and mouse (Abbasi et al., 2021; Driskell et al., 2013; Gay et al., 2020; Guerrero-Juarez et al., 2019; Jiang et al., 2018; Joost et al., 2019; Lim et al., 2018; Rahmani et al., 2014; Rinkevich et al., 2015) skin tissues with important biological relevance in homeostasis, injury-mediated repair and regeneration, disease and cancer. To discern whether cellular and spatial FIB or FIB-like heterogeneity exists in human BCC and PTS regions, we sub-clustered FIB and FIB-like cells based on expression of *PDGFRA* and *RGS5*, yielding a total of 7,080 cells (k_PTS_ = 1305 versus k_BCC_ = 5,775) (**Fig. 3A-B**). Both cell types were collectively positive for extracellular matrix proteins *DCN* and *LUM*. This sub-clustering approach led to the identification of four coarse-grained FIB populations, and two FIB-like populations, all defined by differential expression of unique gene biomarkers (**Fig. 3B-E**).

**Figure 3.**
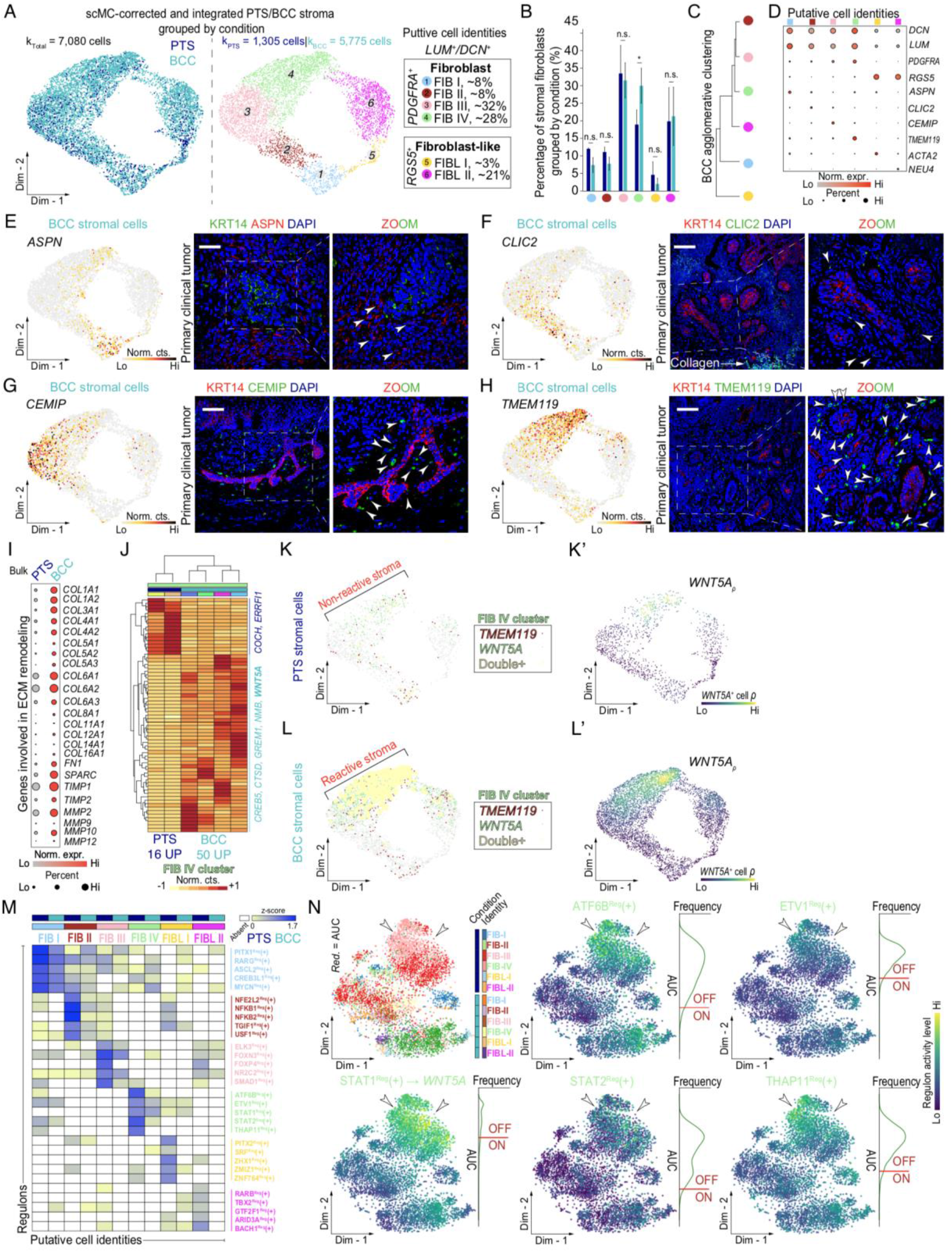
Analysis of stromal cells highlights fibroblast and fibroblast-like cell heterogeneity in human basal cell carcinoma. **A**. Clustering of 7,080 batch-corrected fibroblast/fibroblast-like (FIB/FIB-like) cells from peri-tumor skin (PTS) and basal cell carcinoma (BCC) subtypes grouped by condition. **B**. Clustering of stromal cells grouped by FIB/FIB-like cell subtype. Four putative *PDGFRA*^+^ FIB cell identities were identified, and their identity defined on the right. Two putative *RGS5*^+^ FIB-like cell identities were identified, and their identity defined on the right. **C**. Quantification of FIB and FIB-like cells split by condition. Bar graph is represented as average of cells per donor per cell cluster and bars represent standard error of the mean. Student’s two-tailed t test (\**P* < 0.05, and n.s. – not significant). **D**. Agglomerative clustering of BCC-associated FIB and FIB-like cells. Cells are color-coded as in **B. E**. Dot plots of canonical and marker genes in FIB and FIB-like cell subtypes. Light gray indicates low average gene expression of canonical and marker genes; bright red indicates high average gene expression of canonical and marker genes. Size of circle represents the percentage of cells expressing canonical and marker genes. **E-H**. Feature plots and immunostaining of select BCC-associated FIB and FIB-like cell markers show cluster-specificity and distinct spatial localization in primary clinical tumors. Inset shows magnified area in BCC nest. White arrows point at fibroblasts expressing gene of interest. Sections were counterstained with DAPI. Size bars: 100 µm. **I**. Dotplot demonstrating bulk differential gene expression of genes involved in ECM remodeling in BCC. Light gray indicates low expression of ECM remodeling genes; bright red indicates high expression of ECM remodeling genes. Size of circle represents the percentage of cells expressing ECM remodeling genes. **J**. Heatmap of differentially expressed genes based on normalized counts in peri-tumor and BCC FIB IV. Relevant genes upregulated in BCC FIB IV are highlighted on the right. Bright yellow indicates downregulated genes; dark red indicates upregulated genes. **K, K’, L and L’**. Feature plots show expression of *TMEM119* (dark red), *WNT5A* (light green) or *TMEM119*;*WNT5A* (yellow) in peri-tumor (**K**) versus BCC (**L**) FIBs. Double *TMEM119*;*WNT5A* cells are represented as a bigger circle with respect to *TMEM119*^+^ or *WNT5A*^+^ cells. Feature plots show density of *WNT5A*^+^ cells in peri-tumor (**K’**) versus BCC (**L’**) FIBs. Dark purple indicates low *WNT5A*^+^ cellular density; Bright yellow indicates high *WNT5A*^+^ cellular density. **M**. Heatmap of condition and cell type specific gene regulatory networks demonstrates active regulons in BCC FIB and FIB-like cells compared to peri-tumor skin (Z-score > 0). Light yellow indicates low regulon activity; bright blue indicates high regulon activity; white indicates absent regulon activity. **N**. Regulon activity was used for dimensionality reduction and regulons for BCC FIB IV were visualized in a two-dimensional embedding. White arrows point at BCC FIB IV cluster. Density plots represent AUC distribution per regulon selected. Dark purple indicates low regulon activity; bright indicates high regulon activity. Abbreviations: Norm. expr. – normalized expression; Norm. cts. – normalized counts; AUC – area under the curve.

To explore their gene expression profile and spatial architecture in human BCC, we spatially resolved their distribution *in situ* using protein immunostaining coupled with high-resolution confocal imaging. Cluster 1 fibroblasts (FIB I) represent ∼8% of all FIBs analyzed and collectively express *ASPN* (**Fig. 3F**). ASPN overexpression has been shown to lead to cancer progression and enhanced metastasis, and its expression is similar in mesenchymal stromal cells and cancer-associated fibroblasts (Hughes et al., 2019). *In situ*, ASPN^+^ FIBs appeared ubiquitously throughout the dermis (**Fig. 3F**). Cluster 2 FIBs (FIB II) represent ∼8% of all FIBs and collectively express *CLIC2* (**Fig. 3G**). *In situ*, CLIC2^+^ FIBs are located sparsely surrounding KRT14^+^ tumor cell nests (**Fig. 3G**). Cluster 3 FIBs (FIB III) represented ∼32% of all FIBs and collectively express *CEMIP* (**Fig. 3H**). In colorectal cancer, hypoxia-mediated overexpression of CEMIP in submucosa epithelial cells leads to eventual enhanced cell migration status (Evensen et al., 2015). Additionally, CEMIP^+^ FIBs surround KRT14^+^ tumor cell nests (**Fig. 3H**). Lastly, Cluster 4 FIBs (FIB IV) represent ∼28% of all FIBs and robustly express *TMEM119* (**Fig. 3I**). TMEM119 is upregulated in osteosarcoma cells and its overexpression is associated with increased tumor size, clinical stage, distant metastasis, and poor prognosis (Jiang et al., 2017). The majority of TMEM119^+^ FIBs appeared positioned peripherally and juxtaposed to KRT14^+^ tumor nests (**Fig. 3I**), to a greater extent than those observed for CLIC2^+^ and CEMPI^+^ FIBs or ubiquitous ASPN^+^ FIBs (**Fig. 3F-H**). We also identified two types of *RGS5*^+^ FIB-like cells expressing *ACTA2* (∼3%) and *NEU4* (∼21%). Quantification of cells from each putative FIB and FIB-like subtype partitioned by condition revealed similar cell type frequencies across BCC and PTS samples, with the exception of *TMEM119*^+^ FIBs, which appeared slightly expanded in BCC compared to PTS (**Fig. 3C**). Our *in situ* imaging analysis suggest that TMEM119^+^ FIBs segregate distinctly across KRT14^+^ tumor nests in terms of both position and density, further reinforcing the notion that significant inter-tumoral fibroblast heterogeneity exists in human BCC and that this particular population may be functionally and structurally positioned to support tumoral growth and progression.

We then examined genes coding for ECM-related proteins and compared their expression profiles between conditions to determine the level of ECM remodeling in BCC compared to PTS. In general, we identified prominent changes in extent and expression of genes coding for various collagens, including *COL1A1, COL1A2, COL3A1, COL4A1, COL4A2, COL5A1, COL5A2, COL5A3, COL6A1, COL6A2, COL6A3, COL8A1, COL11A1, COL12A1, COL14A1*, and *COL16A1* (**Fig. 3J**). Analogous to collagen-coding genes, other ECM-related protein-coding genes, including *FN1, SPARC, TIMP1, TIMP2, MMP2, MMP9, MMP10* and *MMP12*, were also enriched in BCC compared to PTS stroma (**Fig. 3J**). The expression of these ECM-related coding genes was not restricted to individual FIB subsets, but rather represent a pan-BCC ECM-related remodeling gene profile. This comparative analysis suggests a large degree of ECM-related remodeling in BCCs compared to PTS that is likely driven by expression of collagen- and metalloproteinase-coding genes.

### Rewiring fibroblasts to a reactive stroma state

Because TMEM119^+^ FIBs segregated distinctly across KRT14^+^ tumor nests and were significantly higher in proportion in BCC samples, we wondered whether they may have a unique gene expression profile that may functionally support tumoral growth and progression. To shed light on this notion, we performed differential gene expression analysis on cluster 4 FIBs across BCC and PTS conditions using a modified version of DEseq2 specifically tailored for single-cell analysis (Love et al., 2014). This analysis led to the identification of sixteen genes differentially upregulated in PTS cluster 4 FIBs, and fifty genes differentially upregulated in BCC cluster 4 TMEM119^+^ FIBs (**Fig. 3K**). One particular gene, *WNT5A*, was overexpressed in BCCs and co-localized with *TMEM119* expression (**Fig. 3L-M**). WNT5A has emerged as an important molecule involved in cancer progression and recent studies have demonstrated that WNT5A regulates cancer cell invasion, metastasis, metabolism, and inflammation (de Sousa and Vermeulen, 2016). Hence, our results suggest a potential functional signaling network of *TMEM119*^+^ FIBs with *KRT14*^+^ tumor cells driven through paracrine non-canonical WNT signaling.

To identify BCC-specific gene regulators (regulons) that may be driving condition-specific gene expression changes in the different FIB populations, including *TMEM119*^+^ FIBs, we performed Gene Regulatory Network (GRN) analysis using pySCENIC (Aibar et al., 2017; Van de Sande et al., 2020) and identified significantly active regulons that were specific to each FIB/FIB-like cluster in BCCs but not in PTS (**Fig. 3N**). The top five regulons active in *TMEM119*^+^ FIBs included ATF6B^REG^(+), ETV1^REG^(+), STAT1^REG^(+), STAT2^REG^(+), and THAP11^REG^(+) (**Fig. 3O**). We found this regulation to be unique across all regulons analyzed. Our single-cell GRN analysis suggests that there are specific regulons that are active in FIB and FIB-like cells, and they differ significantly in activity and regulation of specific targets between BCC and PTS. Furthermore, we found that the STAT1^REG^(+) regulon may be involved in the up-stream regulation of the non-canonical WNT ligand *WNT5A* (**Fig. 3O**).

Our stromal analysis identified a large degree of cellular FIB and FIB-like heterogeneity in human BCC and PTS at gene expression and regulon levels. To determine if these cells exist on a continuum or have distinct cellular states, we calculated the cellular entropy (ξ – energy associated with cellular transitions) of BCC and PTS FIB/FIB-like cells using CEE (Wang et al., 2020) and visualized their individual CEE scores on Waddington landscapes (**Fig. 4A-B**). Our results indicated that BCC FIB populations have lower overall entropy than those of PTS and that *TMEM119*^+^ FIBs show the most stability (**Fig. 5C**), suggesting that FIB I-III may display higher likelihoods of transition to those FIBs that are most juxtaposed to BCC tumor nests (FIB IV). We followed up our analysis with unbiased pseudotime and RNA dynamics-coupled analyses (Bergen et al., 2020a; La Manno et al., 2018; Wang et al., 2019). These complimentary approaches revealed two major, yet distinct trajectories between BCC and PTS. Focusing on FIB cells only, PTS FIBs bifurcated toward *WNT5A*^+^ or *ASPN*^+^ termini from a common *CEMIP*^+^ FIB origin (**Fig. 4D-E**). In sharp contrast, BCC FIBs followed a unilateral trajectory, emanating from *ASPN*^+^ FIBs and culminating in *TMEM119*^+^ FIBs (**Fig. 4F-G**). These observations suggest that a “rewiring” of the tumor stroma takes place to fuel FIBs toward a *TMEM119*^+^/*WNT5A*^+^ state to support tumor growth.

**Figure 4.**
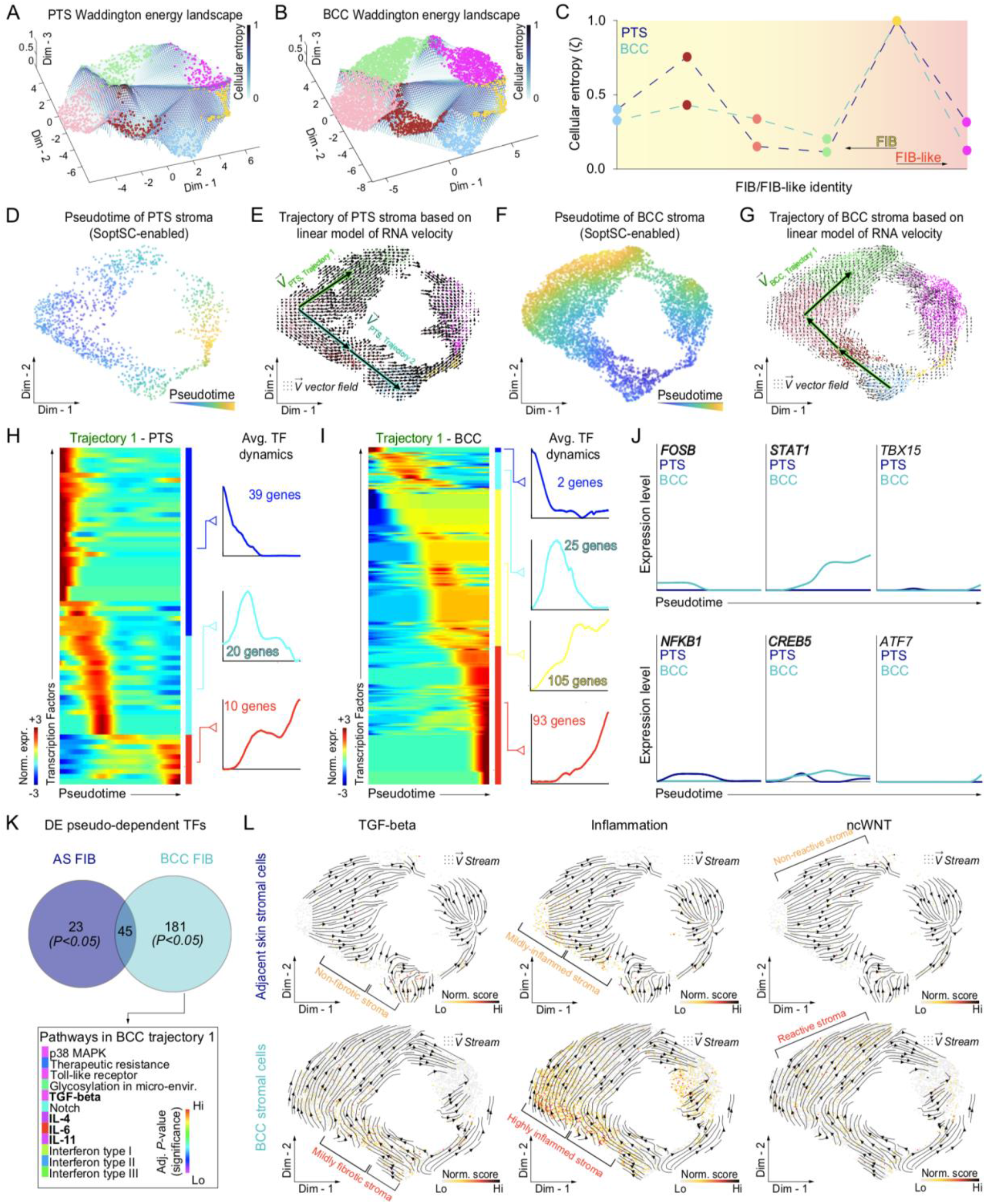
Pseudotemporal and RNA dynamics analyses reveals differential stroma developmental trajectories. **A, B**. Three-dimensional Waddington energy (i.e., entropy) landscape of peri-tumor skin (PTS) and basal cell carcinoma (BCC). Light blue indicates low entropy; dark blue indicates high entropy. **C**. Quantification of cellular energy. Colors of circles correspond to distinct FIB/FIB-like clusters. Dashed lines connect FIB/FIB-like clusters and are color-coded based on type of condition (i.e., PTS versus BCC). **D-G**. Pseudotemporal ordering and RNA velocity-coupled analysis of FIB and FIB-like cells reveal distinct developmental trajectories in PTS and BCC stroma. Cells were scored based on pseudotime and projected on two-dimensional embedding. Dark blue indicates beginning of pseudotime trajectory; dark yellow indicates end of pseudotime trajectory. A linear model of RNA velocity was calculated based on spliced/unspliced ratios. Resultant arrows were projected as vector field on a two-dimensional embedding. Arrows represent direction of cells’ flow. In PTS, bi-directional trajectory of FIBs is represented by trajectory 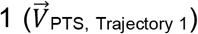 and 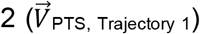. In BCC, unidirection of FIBs is represented by trajectory 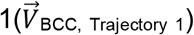. **H, I**. Rolling-wave plots identify pseudo-dependent transcription factors (TFs) overexpressed in PTS (**H**) and BCC (**I**) developmental trajectory 1. Pseudotime levels are based on normalized counts. Dark blue indicates downregulation of TFs; dark red indicates upregulation of TFs. TFs are further grouped depending on their dynamics (k = 3 in PTS; k = 4 in BCC). **J**. Comparison of significant pseudo-dependent transcription factors overexpressed in PTS and BCC developmental trajectories in specific groups. TF dynamics are color-coded based on condition. **K**. Significant pathway ontologies associated with PTS and BCC FIB developmental trajectory 1 (P_adj_ < 0.05). 23 pathway ontologies are unique to PTS; 181 pathway ontologies are unique to BCC. 45 pathway ontologies are shared between PTS and BCC. Specific pathway ontologies in BCC are color-coded based on significance. Rainbow scale: purple indicates low significance; red indicates high significance). **L**. TGF-β, inflammation, and non-canonical WNT pathway scores based on normalized counts overlaid on two-dimensional embedding with linear RNA velocity streams reveal specific pathway programs associated with PTS and BCC stromal developmental trajectory 1. Yellow indicates low score; black indicates high score. Abbreviations: FIB – fibroblasts; DE – differentially expressed; TFs – transcription factors; Norm. score. – normalized score; Norm. cts. – normalized counts.

**Figure 5.**
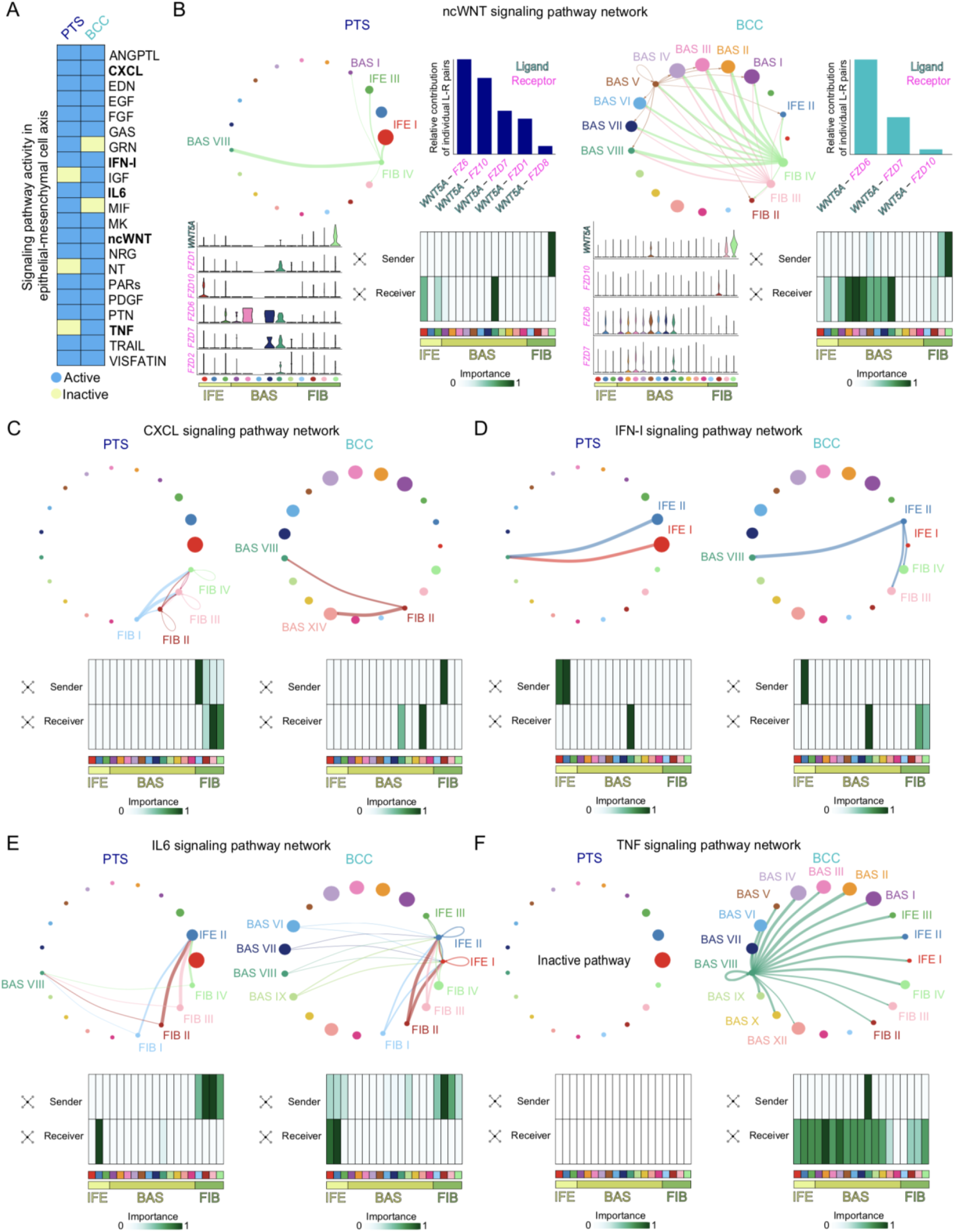
Epithelial-mesenchymal communication modules in basal cell carcinoma delineated by CellChat. **A**. Heatmap of active signaling pathways and their functional relationships of epithelial-mesenchymal crosstalk from peri-tumor skin (PTS) and basal cell carcinoma (BCC) samples. Eighteen signaling pathways are active in PTS and nineteen signaling pathways are active in BCC. Light blue indicates active signaling pathway; light yellow indicates inactive signaling pathway. **B**. ncWNT signaling pathway is differentially active in PTS versus BCC. Circle plots show signaling (i.e., sender) and receiving (i.e., receiver) cells. Color of nodes is congruent with color of senders. Size of cell clusters is representative of the number of active cells in signaling network. FIB-IV has a higher proportion of interactions to epithelial cells via ncWNT in BCC compared to PTS. Bar graphs show distinct relative contribution of specific ligand-receptor pairs for ncWNT signaling in PTS and BCC. *WNT5A* ligand is the only active ligand in the ncWNT signaling network. Violin plots show expression of each ligand and receptor active in the network per cell cluster. Heatmaps describe network centrality analysis showing contribution of ligand (senders) and receptors (receivers). White indicates low importance of network centrality; dark green indicates high importance of network centrality. Importance is on a 0 -1 scale. **C-F**. Circle plots and network centrality analysis for CXCL (**C**), IFN-I (**D**), IL6 (**E**), and TNF (**F**) signaling. Only cell clusters participating in signaling network are labeled. Inactive pathway indicates the pathway is not active in anyone condition. Abbreviations: IFE – interfollicular epidermis; BAS – basal cells; FIB – fibroblasts.

To identify candidate transcription factors (TFs) involved in the acquisition of a *TMEM119*^+^/*WNT5A*^+^ state, we extracted FIBs represented in this trajectory, tree-aligned them in pseudotime with Monocle2 (Qiu et al., 2017), and performed scEpath analysis (Jin et al., 2018) to identify significant, pseudotime-dependent TFs. We identified a total of 69 pseudotime-dependent differentially expressed TFs in PTS 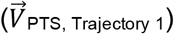 and 225 TFs in BCC 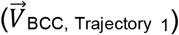 along trajectory 1 (**Fig. 4H-I**). We compared and contrasted TFs from both trajectories by partitioning TFs into groups displaying average TF dynamics, which led to the identification of several genes uniquely present in the BCC trajectory (**Fig. 4J**). Of interest, *STAT1*, and to a lesser extent *TBX15* and *ATF7*, demonstrated pseudo-dependent expression late in the trajectory in BCC compared to PTS toward *TMEM119*^+^ FIBs. Other TFs displayed early pseudo-dependent trajectories and were shut down in *TMEM119*^+^ FIBs, such as *FOSB* in BCC and *NFKB1* in PTS (**Fig. 4J**). To gain a broader view of these TFs and identify major pathways in each trajectory compartment, we performed Gene Ontology (GO) analysis on the pseudotime-dependent TFs (**Fig. 4K**). Among these, we found pathways related to TGF-β, and inflammation to be significantly expressed. We overlaid these GO terms as a biomarker score onto two-dimensional embedding to determine if their expression was closely associated with the rewiring of tumor stroma and overlaid an RNA velocity stream to visualize and match the movement of the cells with their corresponding GOs (**Fig 4L**). Interestingly, *ASPN*^+^ FIBs appear to go through a TGF-β^+^ inflammation state in BCC but not PTS prior to reaching a final reactive stroma status composed of ncWNT signaling-active FIBs – a region high in *WNT5A* ligand. These results suggest that *ASPN*^+^ FIBs become inflamed in BCC and that rewiring of the stroma could arise from inflammation, possibly due to crosstalk with immune cells that have invaded the dermis during BCC progression.

### Inflammatory signaling pathways are highly active in BCC

How FIB state changes influence BCC tumor growth is unclear. To identify signaling differences between the BCC and PTS microenvironments, we probed the human BCC FIB-epithelial interactome by modeling single cell-cell interactions among *KRT14*^+^ epithelial/tumor and FIB/FIB-like cells using CellChat (Jin et al., 2020). We identified 21 significant signaling pathways active in the stroma-epithelial axis, including CXCL, IFN-I, IL6, ncWNT, and TNF (**Fig. 5A**). Although most pathways showed activity in both PTS and BCC, GRN and MIF pathways were inactive in BCC, whereas IGF, NT, and TNF pathways were inactive in PTS (**Fig. 5A**). We then identified differentially regulated signaling pathway ligands and receptors between BCC and PTS by comparing the communication probabilities from cell-cell groups. Unsurprisingly, this approach showed ncWNT as a major signaling pathway highly active in BCC compared to PTS (**Fig. 5B**). The relative contribution of ncWNT signaling was mainly from *WNT5A* ligand to Frizzled receptors. The expression of *WNT5A* ligand in PTS FIBs was minimal and not significant compared to BCC. In sharp contrast, ncWNT signaling was highly active in BCC and mainly driven by *WNT5A* ligand to Frizzled receptors *FZD6, FZD7*, and *FZD10*. The majority of activity was contained in BAS cell clusters as depicted by network centrality analysis of senders and receivers (**Fig. 5B**). WNT5A is a known driver of pro-inflammatory responses, including CXCL, IFN-I, IL6, and TNF (Jung et al., 2013; Pashirzad et al., 2017). CXCL signaling is contained within FIBs in PTS but expands to the BAS cell clusters in BCC (**Fig. 5C**); IFN-I signaling is contained within epithelia in PTS but expands to FIBs in BCC (**Fig. 5D**); IL6 signaling shows greater crosstalk in BCC compared to PTS (**Fig. 5E**); and TNF signaling is exclusive to BCC (**Fig. 5F**). TNF auto- and paracrine signaling originates from cycling epithelial cells in BCC and signals to other epithelial cells and FIBs and is an activator of WNT5A (Zhao et al., 2017). Altogether, these results suggest that acute inflammatory signals may activate WNT5A, which in turn maintains a pro-inflammatory response and acts as a major inflammatory and stress signaling hub center in BCC.

### Heat shock proteins regulate BCC growth

To determine how a pro-inflammatory response from FIBs may influence tumor growth, we looked for relevant differentially expressed genes in BCC versus PTS epithelia and found a significant heat shock protein (HSP) signature in BCC (**Supplementary Fig. 9A**). HSPs are an adaptive response to cellular stress and inflammation and have been strongly implicated in cancer development and progression (Ikwegbue et al., 2019). We identified five HSP-coding genes that were significantly up-regulated in BCC compared to PTS epithelial cells, including *HSP90AB1, HSP90AA1, HSPA2, HSPA6, HSPA8, HSPA12B*, and *HSPA1A*. We next performed GRN analysis using pySCENIC and identified several active regulons that may regulate HSP-coding genes, including ELF1^REG^(+), E2F6^REG^(+), EGR1^REG^(+), EGR2^REG^(+), JUN^REG^(+) and YY1^REG^(+) (**Supplementary Fig. 9B**). These results demonstrate that a high proportion of BCC cells in any one cluster of the distinct BCC subtypes analyzed scored high for HSP gene expression.

We then spatially resolved the expression of HSP70 *in situ* using protein immunostaining coupled with high-resolution confocal imaging as HSP70 is the gene product of HSPA1A. We found that KRT14^+^ BCC nests expressed cytoplasmic HSP70, with seldom IFE epithelial cells and non-epithelial cells expressing the protein (**Fig. 6A-C**). Interestingly, a primary infiltrative BCC with perineural invasion demonstrated nuclear expression of HSP70 (**Fig. 6C**). To determine whether HSPs are important for BCC cell growth, we used the HSP70 inhibitor Ver155008 on the murine BCC cell line ASZ001 and observed a dosage-dependent inhibition of ASZ001 cell proliferation (**Fig. 6D**) with a concomitant decrease in *Gli1* expression, a downstream HH target gene, at both RNA and protein levels (**Fig. 6E-F**). HSP70 inhibition affected both proliferation and survival of BCC cells as determined by Mki67 and Casp3 staining quantification (**Fig. 6G**). Lastly, we aimed to determine the role of HSPs on BCCs *in vivo* using the BCC mouse model *Gli1-Cre*^*ERT2*^; *Ptch1*^*fl/fl*^ (Peterson et al., 2015). We induced *Gli1-Cre*^*ERT2*^; *Ptch1*^*fl/fl*^ mice with tamoxifen for three consecutive days to generate basaloid micro-tumors, followed by intraperitoneal injection with vehicle-control or Ver155008 daily for 7 days (**Fig. 6H**). Histological staining of the dorsal skin of Ver155008-treated mice showed significant reduction in microtumor area compared to vehicle-treated controls in a dose-dependent manner (**Fig. 6I**). Our *in vitro* and *in vivo* studies help to reconcile our scRNA-seq analysis and identify HSPs, particularly HSP70, as a potentially new regulator of BCC tumor growth and may offer a novel therapeutic venue for the treatment of BCCs.

**Figure 6.**
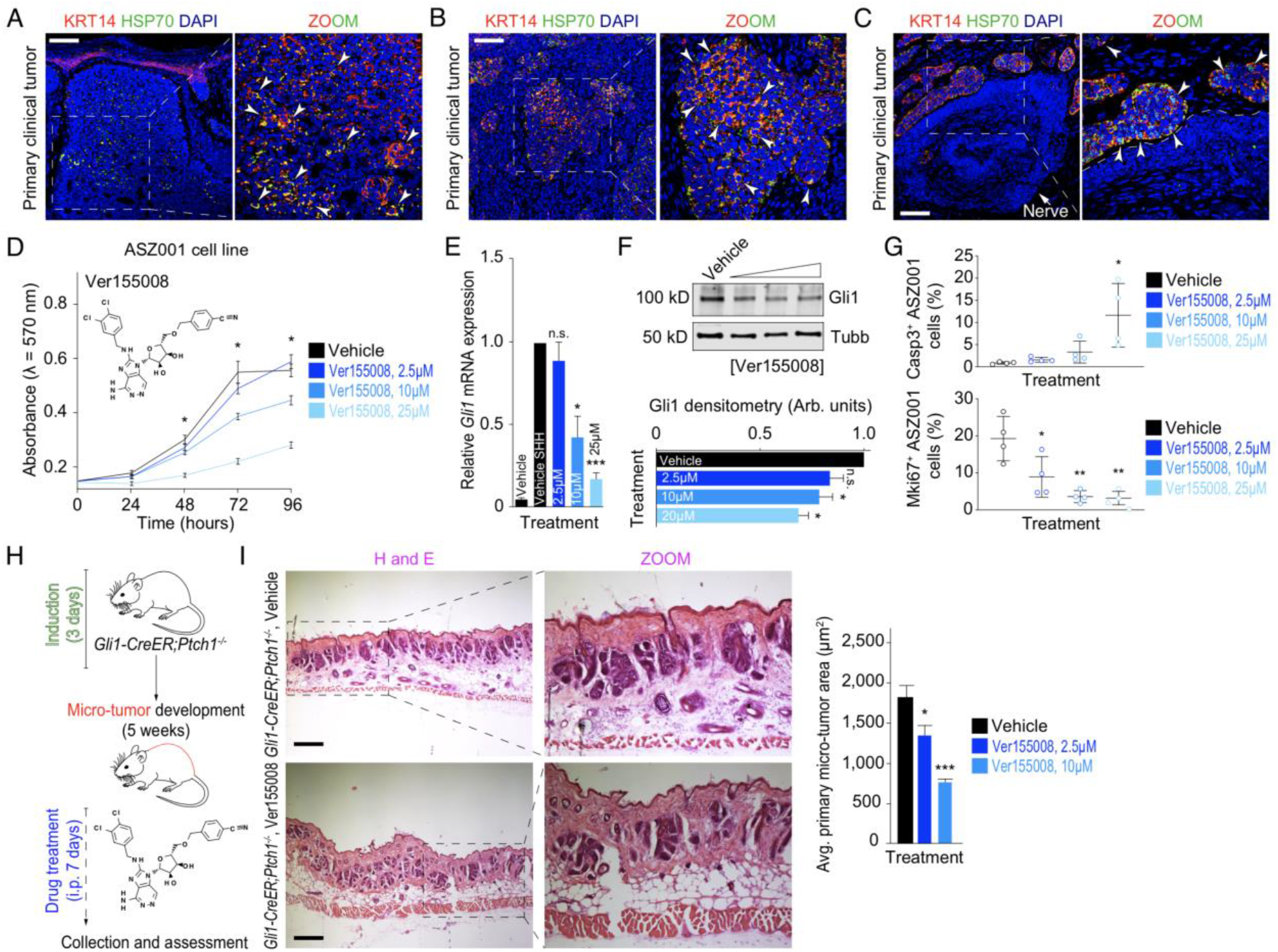
Gene expression profiling and regulon analysis identifies heat shock proteins as prominent regulators of basal cell carcinoma. **A-C**. *In situ* expression of HSP70 protein show distinct spatial localization in primary clinical human basal cell carcinoma (BCC) tumors. Inset shows magnified area in BCC nest. White arrows point at HSP70^+^ epithelial cells in tumor nests. Dashed line demarcates BCC nest from nerve (**C**). Size bars: 100 µm. **D-F**. Heat shock protein (HSP) inhibitor Ver155008 negatively impacts cell growth of ASZ001 murine cells (**D**) and downregulate *Gli1* mRNA (**E**) and protein expression (**F**) *in vitro* in a concentration-dependent manner. Student’s two-tailed t test (\**P* < 0.05, \*\*\**P* < 0.001, and n.s. – not significant). Experiments were repeated at least three times and data is represented as the mean of triplicates ± standard error of the mean (SEM). **G**. HSP inhibitor Ver155008 significantly induces apoptosis via Casp3 and negatively impacts proliferation via Mki67 in ASZ001 murine cells *in vitro* in a concentration-dependent manner. Bar graphs represent the mean of nine replicate wells ± standard error of the mean (SEM). Student’s two-tailed t test (\**P* < 0.05 and \*\**P* < 0.01). **H**. Schematic representation of micro-tumor development and HSP inhibitor treatment in *Gli1-Cre*^*ERT2*^; *Ptch1*^*fl/fl*^ transgenic mice. Ver155008- and vehicle-treated mouse dorsal skin tissues were collected and assessed for micro-tumors. **I**. H&E of Ver155008- and vehicle-treated *Gli1-Cre*^*ERT2*^; *Ptch1*^*fl/fl*^ transgenic mouse dorsal skin tissues. Inset shows magnified area in mouse dermis with localized micro-tumor nests. Size bars: 100 µm. Quantification of micro-tumor surface area in vehicle- and Ver155008-treated *Gli1-Cre*^*ERT2*^; *Ptch1*^*fl/fl*^ transgenic mouse dorsal skin tissues. Surface area decrease in a concentration dependent manner compared to vehicle-treated control. Student’s two-tailed t test (\**P* < 0.05, \*\*\**P* < 0.001).

## Discussion

Functional heterogeneity in human BCC has largely been explored using bulk-level genomic and transcriptomic studies where it was difficult to separate out distinct cell types and clonality within tumors (Atwood et al., 2015; Bonilla et al., 2016; Sharpe et al., 2015). Using single cell technologies, we identified the milieu of cell types and states that make up BCC and found that bulk-level studies can provide complementary datasets but often lead to significant genes that are non-specific and broadly expressed across different cell types due to the heterogeneity of normal and cancerous cells in biopsy samples (**Supplementary Fig. 6-7**). When analyzed at the single cell level, we found additional BCC biomarkers that better define BCCs and label tumors from specific donors that further highlight the heterogeneity of this disease. We also identified spatial heterogeneity in CAFs that led to a trajectory favoring TMEM119^+^/WNT5A^+^ reactive stroma and inflammatory signals that create a burst of cell-cell crosstalk between CAFs and BCC cell clusters. Finally, our results suggest that BCC tumors respond to inflammatory signals from the stroma by expressing HSPs and that HSP inhibitors can be an effective therapeutic to suppress tumor growth.

Our efforts to distinguish between malignant and normal cells between and within biopsy samples to create more nuanced BCC gene signature highlighted the importance of integration benchmarking. Although no algorithm is unflawed (Tran et al., 2020), we demonstrated that use of benchmark integration using several different methods increased confidence in the underlying data. BCCs are highly heterogenous and have the highest mutation frequency out of all cancers (Bonilla et al., 2016), making integration of multiple samples difficult. All four clustering algorithms we used (Seurat, SCTransform, LIGER, Harmony, and scMC) showed remarkable efficiency in correctly clustering non-epithelial cell types with low mutational burden (**Fig. 1C-E, Supplementary Fig. 5**), but epithelial cells with higher mutational burden showed significant batch effects in clustering. In our experience, Seurat, SCTransform, LIGER, and Harmony could not distinguish between normal and malignant cells and Seurat itself clustered each donor separately from each other regardless of origin. However, scMC clustered normal and malignant epithelial cells distinctly while maintaining cohesion within each condition (**Fig. 1D**), likely due to its ability to learn a shared reduced dimensional embedding of cells to retain biological variation while removing technical variation associated with each sample (Zhang and Nie, 2021).

Despite the difficulty in integrating epithelial cells, stromal cell states displayed remarkable cohesion between PTS and BCC samples. Four FIB states and two FIB-like states were found in both normal and malignant samples, suggesting that CAFs may be an active state of normal tissue-resident fibroblasts and that cancer-specific stromal states do not occur in BCC. However, there is a large degree of active remodeling that occurs in BCC stroma, likely driven mainly by collagen and metalloproteinase gene-products (**Fig. 3J**). Furthermore, RNA velocity analysis suggests that highly inflamed stroma expressing TGF-β and IL genes, classic activators of CAFs (Sahai et al., 2020), give rise to reactive stroma highlighted by WNT5A^+^ FIBs (**Fig. 4L**). This cancer-specific rewiring of the stroma goes from an ASPN^+^ state (FIB I) found ubiquitously throughout the stroma, to a CLIC2^+^ (FIB II) and CEMIP^+^ (FIB III) state found sparingly around KRT14^+^ tumor nests, before reaching the TMEM119^+^/WNT5A^+^ state (FIB IV) that surrounds KRT14^+^ tumor nests at a relatively high density compared to the other three FIB states (**Fig. 3F-I**). *TGFB1* and general inflammatory genes are expressed throughout the first three FIB populations and may provide a mechanism of activation to the *WNT5A*^+^ state, while WNT5A may reinforce this signaling as it is a known driver of pro-inflammatory signals to induce an immune response (Jung et al., 2013; Pashirzad et al., 2017). Stromal rewiring driven by inflammation and CAFs are promising therapeutic targets (Sahai et al., 2020) and our GRN analysis suggests the JAK-STAT pathway may regulate *WNT5A* expression, opening up the possibility for JAK-STAT inhibitors in treating BCC patients (Owen et al., 2019).

How CAFs and general FIB inflammation affects BCC tumor growth is unclear. Our CellChat inferred signaling results suggest a burst of signaling between FIBs and BAS clusters involving TNF, IL6, IFN-1, and CXCL pathways (**Fig. 5C-F**). Interestingly, WNT5A is a known driver of each of these pathways (Jung et al., 2013; Pashirzad et al., 2017) and TGFB1 and inflammatory signals like IL6 and TNF are known activators of CAFs and WNT5A in particular (Sahai et al., 2020). With this influx of inflammatory signals, BCCs appear to respond by upregulating HSPs as a protection mechanism (Ikwegbue et al., 2019). HSPs are known to have significant roles in DNA repair mechanisms to maintain genome stability and integrity, a process that is heavily intertwined with inflammation (Techer and Pasero, 2021). Cancers live on a double-edged sword where they need enough genomic instability to thrive but not too much instability to adversely alter successful replication (Andor et al., 2017). Cancer-specific HSP expression may help maintain the genomic instability balance to promote tumor growth, which may explain our results that show HSP inhibitors are effective at suppressing BCC growth. HSP inhibition may be more effective with combinatorial treatment, a likely future direction, as evidenced by on-going clinical trials in cancer types (Mielczarek-Lewandowska et al., 2020).

Overall, our findings illustrate the heterogeneity and dynamic nature of the BCC cellular ecosystem. The signaling relationships between BCC epithelial cells and fibroblasts revealed a WNT5A-inflammatory signature that led to the discovery of a HSP70-specific protective mechanism that is necessary to maintain tumor growth. Further characterizing these types of responses may provide additional mechanistic insight into the complicated crosstalk between the tumor and its microenvironment and provide additional avenues for therapeutic suppression of cancer.

## Experimental procedures and methods

### Ethics statements

Human clinical studies were approved by the Ethics Committee and Institutional Review Board of Stanford University Hospital (Palo Alto, California, USA). We certify that all applicable institutional regulations concerning the ethical use of information and samples from human volunteers were strictly followed in this work. Each subject provided written informed consent. All animal studies were performed in strict adherence to the Institutional Animal Care and Use Committee (IACUC) guidelines of the University of California, Irvine (AUP-21-006).

### Human samples

A total of six surgically discarded tissues (peri-tumor skin, n = 2; basal cell carcinoma, n = 4) were obtained from excisional biopsy specimens at Stanford University Hospital, Palo Alto, California, USA. BCCs were classified into superficial, nodular and infiltrative BCC (ID: BCC-I); superficial and nodular BCC (ID: BCC-II); unknown/”hybrid” (ID: BCC-III); and infiltrative with perineural invasion BCC (ID: BCC-IV) subtypes by board-certified dermatologist/pathologist. All data collection and anonymous analysis was approved by the Institutional Review Board of Stanford University Hospital.

### Mice

*Gli1-Cre*^*ERT2*^; *Ptch1*^*fl/fl*^ mice (Peterson et al., 2015) were genotyped by PCR. Briefly, genomic DNA was collected from mouse toes and lysed in DirectPCR lysis reagent as per manufacturer’s protocol (Fisher Scientific). Genomic DNA was amplified using Taq polymerase master mix (Apex) and products resolved on a 2% agarose gel (Apex). The following primers were used: CreER-Forward: 3’-CATGCTTCATCGTCGGTCC-5’; CreER-Reverse: 3’-GATCATCAGCTACACCAGAG-5’; Patched1-Forward: 3’-AGTGCGTGACACAGATCAGC-5’; and Patched1-Reverse: 3’-CCCAATTACCCATCCTTCCT-5’.

### Micro-tumor induction and drug treatment

Micro-tumors were induced in the skin of six-week-old *Gli1-Cre*^*ERT2*^*;Ptch1*^*fl/fl*^ mice (of indiscriminate gender), by administering 100 μl of 10 mg/ml tamoxifen (Sigma) intraperitoneally for 3 consecutive days. Five weeks later, mice were treated with either DMSO (vehicle) or Ver155008 (16 mg/kg) intraperitoneally for 7 consecutive days. The final volume of all injections was 100 μL. At the end of treatment, mice were sacrificed, and dorsal skin collected, fixed in 4% PFA, immersed in 30% sucrose, and frozen in Tissue Tek OCT compound (Sakura, Japan). Samples were then cryo-sectioned at 14 μm. Unless otherwise noted, at least five mice were used for each treatment condition.

### Micro-tumor assessment

Frozen mouse dorsal skin tissues were cryo-section at 14 μm and stained with H&E (ThermofisherScientific). Images were taken at 200x magnification on an AmScope microscope with an AmScope MU500B digital camera. Micro-tumor size was assessed as the sum of total micro-tumor area and as average size per micro-tumor and quantified using FIJI software (Schindelin et al., 2012). Statistical analysis was performed using GraphPad Prism and application of Student’s two-tailed t test (\**P* < 0.05, \*\**P* < 0.01, \*\*\**P* < 0.001, and n.s. – not significant).

### Histology and immunohistochemistry

Discarded human tumor skin tissues were processed at the Department of Pathology at Stanford University and sectioned at a thickness of 5 μm. Immunostaining was performed on paraffin sections. Heat-based antigen retrieval was performed when necessary. Tissue sections were blocked in either 3% BSA or 3% Donkey serum. The following primary antibodies were used: rabbit anti-ASPN (Abcam, ab58741, 1:100), rabbit anti-CLIC2 (Abcam, ab175230, 1:50), rabbit anti-CEMIP (Proteintech, 211291AP, 1:100), rabbit anti-TMEM119 (Proteintech, 27585-1-AP, 1:100), chicken anti-KRT14 (BioLegend, SIG-3476, 1:1000), mouse anti-LHX2 (Santa Cruz Biotechnology, sc-517243, 1:50), mouse anti-ACTA2 (R&D Systems, MAB1420, 1:500), mouse anti-KRT15 (Santa Cruz Biotechnology, sc-47697, 1:1000), rabbit anti-HSP70 (Proteintech, 10995-1-AP, 1:1000), rabbit anti-CASP3 (R&D Systems, MAB835-SP, 1:1000), and rabbit anti-MKI67 (Abcam, ab15580, 1:1000). Secondary antibodies were used at a concentration of 1:1000. Sections were counterstained with DAPI (VectorLabs). Images were acquired on an Olympus FV3000 confocal laser scanning microscope (Germany).

### Cell culture and growth assay

ASZ001 cells were grown in 154CF media containing chelated 2% FBS (Life Technologies), 1% Penicillin-streptomycin (Life Technologies), and 0.07 mM CaCl_2_ (Life Technologies). NIH 3T3 cells were grown in DMEM (Life Technologies) containing 10% FBS (Life Technologies) and 1% Penicillin-streptomycin (Life Technologies) and were incubated in a water-jacketed incubator at 37°C with 5% CO_2_ output. ASZ001 cells were seeded at a density of 1,000 cells/well into 96-well flat-bottom plates. After 24 hours, cells were treated with DMSO (vehicle control) or varying concentrations of Ver155008 (MedChemExpress) consecutively for 2, 4, and 6 days. Growth assay was performed with MTT (Sigma-Aldrich) per manufacturer’s protocol. Proliferation (MKI67) and apoptosis (CASP3) were determined by immunostaining fixed cells at the indicated time points. Unless otherwise noted, experiments were repeated at least three times and data is represented as the mean of nine replicate wells ± standard error of the mean (SEM). Statistical analysis was performed using GraphPad Prism and application of Student’s two-tailed t test (\**P* < 0.05, \*\**P* < 0.01, and \*\*\**P* < 0.001).

### Hedgehog assay

NIH 3T3 cells were seeded to confluence, serum-starved (SS) or serum-starved in 1:100 SHH-N ligand (SSH) with DMSO (vehicle control) or Ver155008 (MedChemExpress) at various concentrations for 24 hours. RNA was isolated using the Direct-zol RNA MiniPrep Plus (ZYMO Research). Quantitative RT-PCR was performed using the iTaq Univer SYBR Green 1-Step Kit (Bio-Rad) on a StepOnePlus Real-time PCR system (Applied BioSystem). The fold change in mRNA expression of the HH target gene *Gli1* was measured using ΔΔC_t_ analysis with *Gapdh* as an internal control gene. Unless otherwise noted, experiments were repeated at least three times and data is represented as the mean of triplicates ± standard error of the mean (SEM). Statistical analysis was performed using GraphPad Prism and application of Student’s two-tailed t test (\**P* < 0.05, \*\**P* < 0.01, \*\*\**P* < 0.001, and n.s. – not significant).

### Cell isolation and 3’-droplet-enabled single-cell RNA-sequencing

Adjacent peri-tumor and tumor skin specimens were surgically excised from human donors at Stanford University Hospital and immediately shipped to University of California, Irvine. Within 24 hours, excised tissues were minced and incubated in a Dispase II (Sigma-Aldrich) and Collagenase IV (Sigma-Aldrich) solution overnight at 4°C. Cells were incubated in 0.25% Trypsin-EDTA for 15 min at 37 °C and quenched with chelated FBS. Cells were passed through a 40 μm filter, centrifuged at 1500 rpm for 5 min, and the pellet resuspended in Keratinocyte Serum Free Media supplemented with Epidermal Growth Factor 1-53 and Bovine Pituitary Extract (Life Technologies; 17005042). Following isolation, cells were resuspended in PBS free of Ca_2_^+^ and Mg_2_^+^ and 1% BSA and stained with SYTOX Blue Dead Cell Stain (ThermoFisher; S34857). Samples were bulk sorted at 4°C on a BD FACSAria Fusion using a 100 μm nozzle (20 PSI) at a flow rate of 2.0 with a maximum threshold of 3000 events/s. Following exclusion of debris and singlet/doublet discrimination, cells were gated on viability for downstream scRNA-seq. Live cells were resuspended in 0.04% UltraPure BSA (Sigma-Aldrich) and counted using the automated cell counter Countess (Thermo). GEM generation, barcoding, post GEM-RT cleanup, cDNA amplification and cDNA library construction were performed using Single Cell 3’ v2 chemistry (10X Genomics). cDNA libraries were sequenced on an Illumina HiSeq4000 platform (Illumina) (one lane, 100 PE). Cell counting, suspension, GEM generation, barcoding, post GEM-RT cleanup, cDNA amplification, library preparation, quality control, and sequencing were performed at the Genomics High Throughput Sequencing facility at the University of California, Irvine.

### 3’-droplet-enabled single-cell RNA-sequencing raw data processing

Transcripts were aligned to the human reference genome (GRCH38/transcriptome) using CellRanger (version 2.1.0). Sequencing metrics for each library are as follow: (PTS-I) <u>Sequencing metrics</u>: ∼264,949,873 total number of reads, ∼98.7% valid barcodes; <u>Mapping metrics</u>: ∼93.1% reads mapped to genome, ∼91.0% reads mapped confidently to genome, ∼71.1% reads mapped confidently to transcriptome; <u>Cell metrics</u>: ∼7,164 estimated number of cells, ∼92.9% fraction reads in cells, ∼36,983 mean reads per cell, ∼2,382 median genes per cell, ∼21,853 total genes detected, ∼9,238 median UMI counts per cell. (PTS-II) <u>Sequencing metrics</u>: ∼317,022,706 total number of reads, ∼98.7% valid barcodes; <u>Mapping metrics</u>: ∼93.1% reads mapped to genome, ∼91.0% reads mapped confidently to genome, ∼71.1% reads mapped confidently to transcriptome; <u>Cell metrics</u>: ∼7,164 estimated number of cells, ∼92.9% fraction reads in cells, ∼36,983 mean reads per cell, ∼2,382 median genes per cell, ∼21,853 total genes detected, ∼9,238 median UMI counts per cell. (BCC-I) <u>Sequencing metrics</u>: ∼170,434,662 total number of reads, ∼98.5% valid barcodes; <u>Mapping metrics</u>: ∼91.3% reads mapped to genome, ∼89.0% reads mapped confidently to genome, ∼67.6% reads mapped confidently to transcriptome; <u>Cell metrics</u>: ∼10,025 estimated number of cells, ∼90.2% fraction reads in cells, ∼17,000 mean reads per cell, ∼2,484 median genes per cell, ∼22,986 total genes detected, ∼6,618 median UMI counts per cell. (BCC-II) <u>Sequencing metrics</u>: ∼128,178,058 total number of reads, ∼98.5% valid barcodes; <u>Mapping metrics</u>: ∼92.8% reads mapped to genome, ∼90.6% reads mapped confidently to genome, ∼70.4% reads mapped confidently to transcriptome; <u>Cell metrics</u>: ∼12,487 estimated number of cells, ∼86.8% fraction reads in cells, ∼10,264 mean reads per cell, ∼1,708 median genes per cell, ∼22,737 total genes detected, ∼4,361 median UMI counts per cell. (BCC-III) <u>Sequencing metrics</u>: ∼335,812,707 total number of reads, ∼98.3% valid barcodes; <u>Mapping metrics</u>: ∼87.5% reads mapped to genome, ∼84.8% reads mapped confidently to genome, ∼65.7% reads mapped confidently to transcriptome; <u>Cell metrics</u>: ∼7,094 estimated number of cells, ∼82.6% fraction reads in cells, ∼47,337 mean reads per cell, ∼2,315 median genes per cell, ∼23,364 total genes detected, ∼8,292 median UMI counts per cell. (BCC-IV) <u>Sequencing metrics</u>: ∼277,281,459 total number of reads, ∼98.6% valid barcodes; <u>Mapping metrics</u>: ∼93.2% reads mapped to genome, ∼90.6% reads mapped confidently to genome, ∼62.6% reads mapped confidently to transcriptome; <u>Cell metrics</u>: ∼8,829 estimated number of cells, ∼88.8% fraction reads in cells, ∼31,405 mean reads per cell, ∼1,983 median genes per cell, ∼23,362 total genes detected, ∼5,516 median UMI counts per cell.

### Doublet/multiplet simulation and low-quality cell pruning

Putative doublets/multiplets were simulated with Single-Cell Remover of Doublets (Scrublet) (version 0.2.1) (Wolock et al., 2019) using raw count matrices. The number of neighbors used to construct the KNN classifier of observed transcriptomes and simulated doublets/multiplets was set as default. The doublet/multiplet score threshold was adjusted manually as suggested by the developer. Briefly, digital matrices for putative singlets were used for low-quality cell pruning using a user-defined pipeline. Viable singlets were kept and used for downstream query and comparative analyses if and only if they met the following collective quality control criteria: a) 350 < genes/cell < 5000; b) cells contained no more than 10% of mitochondrial gene expression; c) cells were not identified as outliers (P-value 1e-3) (Fan et al., 2016).

### Data processing and benchmarking of 3’-droplet-enabled single-cell RNA-sequencing

*<u>Processing of individual data sets</u>*. Pre-processed digital matrices from individual tumor data sets were processed using Seurat (version 4.0.1). Seurat objects were created and Log-normalized with a scale factor of 10,000. Variable features were identified using *vst* with top 2,000 features. Data was scaled and metadata variables, including counts and percent mitochondria, were regressed. PCA was calculated using variable features identified using a combination of heuristic and statistical approaches. Individual data sets were visualized using a two-dimensional embedding. *<u>Benchmarking of integrated data sets</u>*. Individually processed Seurat objects used for integration, downstream analyses, or visualization with Seurat (Version 3.0.0900) (Stuart et al., 2019), Single Cell Transform (version 0.3.2) (Hafemeister and Satija, 2019), LIGER (Version 2.0.1) (Welch et al., 2019), Harmony (Version 0.1.0) (Korsunsky et al., 2019), or scMC (Version 1.0.0) (Zhang and Nie, 2021).

### Copy number variation analysis

To infer genomic copy number structure, we performed InferCNV (version 1.7.1) as per developer’s suggestions using standard parameters (https://github.com/broadinstitute/inferCNV/wiki). We used a cutoff of 0.1 for the minimum average read counts per gene among reference cells. We used non-immune, non-epithelial, and non-appendage epithelial cells as internal reference control.

### Gene marker analysis

To identify shared (i.e., general) and unique (i.e., subtype/donor-specific) BCC gene signature profiles in our dataset, we performed differential gene expression analysis (i.e., gene marker identification) using Seurat (version 4.0.1). For this, we implemented a minimum average Log_2_ fold change in expression (0.25x) and a minimum percent of cells that must express gene marker in either one cluster (0.50 or 50%). Gene markers were identified using the *FindAllMarkers* function using the Wilcoxon Rank Sum Test.

### Marker gene module scoring

Aggregate marker gene module scores were assigned to early epithelial differentiation, late epithelial differentiation, and BCC identity score using the AddModuleScore function in Seurat with the following conditions enabled: ctrl = 2.5 or 5. Basal, non-differentiating and basal late differentiating epithelial cells were identified by implementation of a “Early differentiation” and “Late differentiation” signatures defined by a core set of known markers as previously described (Lopez-Pajares et al., 2015). BCC cells were identified by implementation of a BCC signature defined by a core set of known markers as previously described (Yost et al., 2019). Hedhehog- and WNT-active and responsive cells were identified by implementation of a “Hedgehog signaling aggregate score” or “WNT signaling aggregate score” defined by a core set of known HH and WNT ligands and receptors, respectively as previously described in GSEA (Subramanian et al., 2005). Hedhehog and WNT-active/responsive cells were colored distinctly. Double-positive cells were color-coded based on a blend threshold score. Aggregate marker gene module and blend threshold scores were Log-normalized and visualized in two-dimensional feature plots.

### Quantification of cells

To quantify cells expressing our genes of interest, we instituted the following approach: *Gene A*^+^ (i.e., *BCAM*) represents a group of cells whose gene expression for “Gene A” per cell in a group (i.e., condition) exceeds the maximum expression for “Gene A” among all cells multiplied by a constant of 0.8. Such notation is also applied to other genes of interest (i.e., *EPCAM, TP63, LGALS1, MYLK, CALML5, SCGB2A2*, or *KRT19*). If a cell belongs to the cluster *BCAM*^+^, *EPCAM*^+^, *TP63*^+^, and *LGALS1*^+^ then that particular cell will be clustered as “Quadrupled^+^”. For all the other cases, the cell is clustered as “None”. Calculations were carried out in MATLAB (Version 9.5).

### Visualization of single-cell RNA-sequencing data

*Cell density plots*. Cell density plots were calculated using Nebulosa (version 1.0.2) and overlaid on a two-dimensional embedding (Alquicira-Hernandez and Powell, 2021).

### Differential gene expression analysis

Differentially expressed genes between adjacent and tumor skin fibroblasts were calculated using DESeq2 as previously described and with minor modifications tailored for the analysis of scRNA-seq data sets (Love et al., 2014). Hypothesis testing was performed with the Wald Test and differentially expressed genes for a particular comparison (i.e., adjacent peri-tumor versus tumor from anyone cluster) were filtered using a P-adjusted threshold of 0.05 and a fold-change of 1.5x in either direction.

### Gene ontology analysis

Gene Ontology (GO) analysis was performed with Panther (Thomas et al., 2003), DAVID (Huang et al., 2007), and Enrichr (Chen et al., 2013; Kuleshov et al., 2016). GOs were visualized by heatmap.

### RNA dynamics analysis

To calculate RNA dynamics in single cells we performed RNA velocity analysis. First, we generated loom files using the Python script velocyto.py (Python version 2.7.2) for each individual library. Loom files for individual libraries from a particular condition (normal or tumor skin) were combined using loompy (version 2.0.16). Velocity vectors were estimated using scVelo (version 0.2.2) based on the linear or dynamic model of RNA velocity with default parameters (Bergen et al., 2020b). Velocity vectors were overlaid on a two-dimensional embedding. Predicted root and terminal states based on Markovnikov reaction diffusion were calculated. Root and terminal states were visualized on a two-dimensional embedding.

### Pseudotime analyses

Pseudo-ordering of individual fibroblasts (FIBs) from peri-tumor skin (PTS) or basal cell carcinoma (BCC) was performed using Monocle2 (Version 2.10.1) (Qiu et al., 2017; Trapnell et al., 2014). Briefly, FIB cells from PTS or BCC trajectory 1 were subclustered and a *cellDataSet* object was created in Monocle2 with the function *newCellDataSet* with the following arguments enabled: *lowerDetectionLimit = 1*.*0, expressionFamily = negbinomial*.*size()*. Subclustered FIBs were ordered based on variable features and DDRTree-based dimensionality reduction was performed using the *reduceDimensions* function. To identify differentially expressed pseudotime-dependent transcription factor changes, we applied single cell Energy path (scEpath; Version 1; MATLAB Version 9.5) (Jin et al., 2018) on Monocle2-ordered PTS or BCC FIBs. Statistically significant pseudotime-dependent gene changes were identified by comparing the standard deviation of the observed smoothed expressions with a set of similarly permuted expressions by randomly permuting the cell order (*nboot* = 100 permutations). We considered all genes with a standard deviation greater than 0.01 and a Bonferroni-corrected P-value below a significance level *α* = 0.05 to be pseudotime-dependent. Human transcription factors were identified using the Animal Transcription Factor Database (AnimalTFDB 2.0) (Zhang et al., 2015) by enabling the *TF_Ifo*.*human*.*Symbol* function. Pseudotime-dependent genes were represented and visualized using a rolling wave plot with user-defined optimal K-means clustering.

### Cellular entropy estimation

Cellular entropy estimation was performed as previously described with minor modifications and projected on a three-dimensional Waddington energy landscape and quantified (Wang et al., 2020).

### Cell-cell communication analysis

Cell-cell communication networks were modeled based on abundance of ligand-receptor pair transcripts with CellChat (version 0.5.0) (Jin et al., 2021). To infer cell-cell communication network differences and similarities between adjacent and tumor cells, we sub-clustered epithelial and fibroblasts from individual conditions. Cell groups of interest were merged, normalized and used as input for CellChat. We calculated over-expressed genes and the significant ligand-receptor interactions with the *identifyOverExpressedGenes* (thresh.p = 0.05) and *identifyOverExpressedInteractions* functions, respectively. We used the human database of ligand-receptor pairs provided by CellChat. The communication probabilities were calculated with *computeCommunProb* (tresh = 0.05, nboot = 100, Hill function parameter kn = 0.5) and inferred the cellular communication network at a signaling pathway level using *computeCommunProbPathway* with default parameters (tresh = 0.05). We filtered out cell-cell communications where a minimum of 5 cells per group were present. The aggregated cell-cell communication networks were calculated with *aggregateNet* (tresh = 0.05) with default parameters. To identify conserved and induced/perturbed communication networks in adjacent and tumor data sets, we performed joint manifold and classification learning analyses.

### Gene Regulatory Network analysis

Gene Regulatory Networks were modeled with pySCENIC (Aibar et al., 2017; Van de Sande et al., 2020) (version 0.10.2) in a Python Environment (version 3.7). We used a pre-defined list of human transcription factors (TFs) (https://github.com/aertslab/pySCENIC/blob/master/resources/hs_hgnc_tfs.txt) and inferred regulatory interactions between them and their putative target genes via GRNBoost2. We focused on activating modules and used them for downstream query. We performed cisTarget motif enrichment based on a ranking and recovery approach with a 10kb putative regulatory region boundary from the TSS (hg38_refseq-r80_10kb_up_and_down_tss.mc9nr.feather) (https://resources.aertslab.org/cistarget/). AUC metrics were used to assess gene recovery and regulon activity and used for dimensionality reduction. The activity of regulons across fibroblast/fibroblast-like cells were compared against each other and across conditions by converting Regulon Specificity Scores (RSS) to Z-scores as previously described. Regulon-specific modules were identified from the network inference output using iRegulon (http://iregulon.aertslab.org). Regulon targets were correlated with differentially expressed genes in specific clusters across conditions.

### Data repository and code availability

The authors declare that all supporting data are available within the Article and its Supplementary Information files. Raw scRNA-seq matrices have been deposited in the Gene Expression Omnibus (GEO) database under the accession code: GSE141526. All generic and custom R, Python, MATLAB, and shell scripts are available on reasonable request.

## Acknowledgements

S.X.A. is supported by NIH grant R01CA237563 and the Concern Foundation (CF204525). K.Y.S. is supported by NIH Grant 5K23CA211793 and is the D. G. “Mitch” Mitchell Clinical Investigator supported by the Damon Runyon Cancer Research Foundation (CI-104-19). Q.N. is supported by NIH grants R01GM123731 and U01AR07315, NSF grant DMS11763272, a grant by Jayne Koskinas Ted Giovanis Foundation for Health and Policy jointly with the Breast Cancer Research Foundation, and an NSF-Simons Foundation grant (594598). C.F.G-J. is supported by UC Irvine Chancellor’s ADVANCE Postdoctoral Fellowship Program, NSF-Simons Postdoctoral Fellowship, and NSF Grant DMS1763272 (Q.N.) and a kind gift from the Howard Hughes Medical Institute Hanna H. Gray Postdoctoral Fellowship Program. This project was partly funded by the UCI Center for Complex Biological Systems (CCBS) opportunity award and the UCI Office of Research. The authors wish to acknowledge the support of the Chao Family Comprehensive Cancer Center Optical Biology Core Shared Resource, supported by the National Cancer Institute of the NIH under award number P30CA062203. We thank Jennifer Atwood and the UCI Institute for Immunology Flow Cytometry Core Facility for help with cell sorting.

## Author contributions

S.X.A., C.F.G.-J., G.H.L., and K.S. conceived the project; S.X.A., Q.N., and K.S. supervised research; K.S. and S.A. acquired peri-tumor and tumor clinical samples; M.L.D. isolated single cells for library preparation; C.F.G-J., Y.L., G.H.L., and S.W. performed bioinformatic and scRNA-seq data analyses; C.F.G-J. curated single-cell RNA-sequencing data; Y.L. performed confocal imaging experiments; C.F.G-J., Y.L., T.T.L.N., R.Y.C., Y.S., and M.K. performed research; C.F.G-J., G.H.L., K.S., and S.X.A. analyzed and interpreted data; C.F.G-J. produced figures with input from G.H.L.; C.F.G-J., G.H.L., and S.X.A. wrote the manuscript; all authors discussed the results and commented on the manuscript.

## REFERENCES

Abbas, O., Richards, J.E., and Mahalingam, M. (2010). Fibroblast-activation protein: a single marker that confidently differentiates morpheaform/infiltrative basal cell carcinoma from desmoplastic trichoepithelioma. Mod Pathol 23, 1535–1543.

Abbasi, S., Sinha, S., Labit, E., Rosin, N.L., Yoon, G., Rahmani, W., Jaffer, A., Sharma, N., Hagner, A., Shah, P., et al. (2021). Distinct Regulatory Programs Control the Latent Regenerative Potential of Dermal Fibroblasts during Wound Healing. Cell Stem Cell 28, 581–583.

Adegboyega, P.A., Rodriguez, S., and McLarty, J. (2010). Stromal expression of actin is a marker of aggressiveness in basal cell carcinoma. Hum Pathol 41, 1128–1137.

Aibar, S., Gonzalez-Blas, C.B., Moerman, T., Huynh-Thu, V.A., Imrichova, H., Hulselmans, G., Rambow, F., Marine, J.C., Geurts, P., Aerts, J., et al. (2017). SCENIC: single-cell regulatory network inference and clustering. Nat Methods 14, 1083–1086.

Alquicira-Hernandez, J., and Powell, J.E. (2021). Nebulosa recovers single cell gene expression signals by kernel density estimation. Bioinformatics.

Andor, N., Maley, C.C., and Ji, H.P. (2017). Genomic Instability in Cancer: Teetering on the Limit of Tolerance. Cancer Res 77, 2179–2185.

Atwood, S.X., Li, M., Lee, A., Tang, J.Y., and Oro, A.E. (2013). GLI activation by atypical protein kinase C iota/lambda regulates the growth of basal cell carcinomas. Nature 494, 484–488.

Atwood, S.X., Sarin, K.Y., Whitson, R.J., Li, J.R., Kim, G., Rezaee, M., Ally, M.S., Kim, J., Yao, C., Chang, A.L., et al. (2015). Smoothened variants explain the majority of drug resistance in basal cell carcinoma. Cancer Cell 27, 342–353.

Bakshi, A., Chaudhary, S.C., Rana, M., Elmets, C.A., and Athar, M. (2017). Basal cell carcinoma pathogenesis and therapy involving hedgehog signaling and beyond. Mol Carcinog 56, 2543–2557.

Basset-Seguin, N., Sharpe, H.J., and de Sauvage, F.J. (2015). Efficacy of Hedgehog pathway inhibitors in Basal cell carcinoma. Mol Cancer Ther 14, 633–641.

Bergen, V., Lange, M., Peidli, S., Wolf, F.A., and Theis, F.J. (2020a). Generalizing RNA velocity to transient cell states through dynamical modeling. Nat Biotechnol.

Bergen, V., Lange, M., Peidli, S., Wolf, F.A., and Theis, F.J. (2020b). Generalizing RNA velocity to transient cell states through dynamical modeling. Nat Biotechnol 38, 1408–1414.

Bernemann, T.M., Podda, M., Wolter, M., and Boehncke, W.H. (2000). Expression of the basal cell adhesion molecule (B-CAM) in normal and diseased human skin. J Cutan Pathol 27, 108–111.

Bhowmick, N.A., Neilson, E.G., and Moses, H.L. (2004). Stromal fibroblasts in cancer initiation and progression. Nature 432, 332–337.

Bircan, S., Candir, O., Kapucoglu, N., and Baspinar, S. (2006). The expression of p63 in basal cell carcinomas and association with histological differentiation. J Cutan Pathol 33, 293–298.

Bolander, A., Agnarsdottir, M., Stromberg, S., Ponten, F., Hesselius, P., Uhlen, M., and Bergqvist, M. (2008). The protein expression of TRP-1 and galectin-1 in cutaneous malignant melanomas. Cancer Genomics Proteomics 5, 293–300.

Bonilla, X., Parmentier, L., King, B., Bezrukov, F., Kaya, G., Zoete, V., Seplyarskiy, V.B., Sharpe, H.J., McKee, T., Letourneau, A., et al. (2016). Genomic analysis identifies new drivers and progression pathways in skin basal cell carcinoma. Nat Genet 48, 398–406.

Cameron, M.C., Lee, E., Hibler, B.P., Barker, C.A., Mori, S., Cordova, M., Nehal, K.S., and Rossi, A.M. (2019a). Basal cell carcinoma: Epidemiology; pathophysiology; clinical and histological subtypes; and disease associations. J Am Acad Dermatol 80, 303–317.

Cameron, M.C., Lee, E., Hibler, B.P., Giordano, C.N., Barker, C.A., Mori, S., Cordova, M., Nehal, K.S., and Rossi, A.M. (2019b). Basal cell carcinoma: Contemporary approaches to diagnosis, treatment, and prevention. J Am Acad Dermatol 80, 321–339.

Chahal, H.S., Wu, W., Ransohoff, K.J., Yang, L., Hedlin, H., Desai, M., Lin, Y., Dai, H.J., Qureshi, A.A., Li, W.Q., et al. (2016). Genome-wide association study identifies 14 novel risk alleles associated with basal cell carcinoma. Nat Commun 7, 12510.

Chang, A.L., and Oro, A.E. (2012). Initial assessment of tumor regrowth after vismodegib in advanced Basal cell carcinoma. Arch Dermatol 148, 1324–1325.

Chang, A.L., Solomon, J.A., Hainsworth, J.D., Goldberg, L., McKenna, E., Day, B.M., Chen, D.M., and Weiss, G.J. (2014). Expanded access study of patients with advanced basal cell carcinoma treated with the Hedgehog pathway inhibitor, vismodegib. J Am Acad Dermatol 70, 60–69.

Chen, E.Y., Tan, C.M., Kou, Y., Duan, Q., Wang, Z., Meirelles, G.V., Clark, N.R., and Ma’ayan, A. (2013). Enrichr: interactive and collaborative HTML5 gene list enrichment analysis tool. BMC Bioinformatics 14, 128.

Chetry, M., Song, Y., Pan, C., Li, R., Zhang, J., and Zhu, X. (2020). Effects of Galectin-1 on Biological Behavior in Cervical Cancer. J Cancer 11, 1584–1595.

Chow, R.Y., Jeon, U.S., Levee, T.M., Kaur, G., Cedeno, D.P., Doan, L.T., and Atwood, S.X. (2021a). PI3K Promotes Basal Cell Carcinoma Growth Through Kinase-Induced p21 Degradation. Front Oncol 11, 668247.

Chow, R.Y., Levee, T.M., Kaur, G., Cedeno, D.P., Doan, L.T., and Atwood, S.X. (2021b). MTOR promotes basal cell carcinoma growth through atypical PKC. Exp Dermatol 30, 358–366.

Clark, C.M., Furniss, M., and Mackay-Wiggan, J.M. (2014). Basal cell carcinoma: an evidence-based treatment update. Am J Clin Dermatol 15, 197–216.

de Sousa, E.M.F., and Vermeulen, L. (2016). Wnt Signaling in Cancer Stem Cell Biology. Cancers (Basel) 8.

de Vries, E., Louwman, M., Bastiaens, M., de Gruijl, F., and Coebergh, J.W. (2004). Rapid and continuous increases in incidence rates of basal cell carcinoma in the southeast Netherlands since 1973. J Invest Dermatol 123, 634–638.

Depianto, D., Kerns, M.L., Dlugosz, A.A., and Coulombe, P.A. (2010). Keratin 17 promotes epithelial proliferation and tumor growth by polarizing the immune response in skin. Nat Genet 42, 910–914.

Driskell, R.R., Lichtenberger, B.M., Hoste, E., Kretzschmar, K., Simons, B.D., Charalambous, M., Ferron, S.R., Herault, Y., Pavlovic, G., Ferguson-Smith, A.C., et al. (2013). Distinct fibroblast lineages determine dermal architecture in skin development and repair. Nature 504, 277–281.

Eberl, M., Mangelberger, D., Swanson, J.B., Verhaegen, M.E., Harms, P.W., Frohm, M.L., Dlugosz, A.A., and Wong, S.Y. (2018). Tumor Architecture and Notch Signaling Modulate Drug Response in Basal Cell Carcinoma. Cancer Cell 33, 229–243 e224.

Evensen, N.A., Li, Y., Kuscu, C., Liu, J., Cathcart, J., Banach, A., Zhang, Q., Li, E., Joshi, S., Yang, J., et al. (2015). Hypoxia promotes colon cancer dissemination through up-regulation of cell migration-inducing protein (CEMIP). Oncotarget 6, 20723–20739.

Fan, J., Salathia, N., Liu, R., Kaeser, G.E., Yung, Y.C., Herman, J.L., Kaper, F., Fan, J.B., Zhang, K., Chun, J., et al. (2016). Characterizing transcriptional heterogeneity through pathway and gene set overdispersion analysis. Nat Methods 13, 241–244.

Gao, R., Bai, S., Henderson, Y.C., Lin, Y., Schalck, A., Yan, Y., Kumar, T., Hu, M., Sei, E., Davis, A., et al. (2021). Delineating copy number and clonal substructure in human tumors from single-cell transcriptomes. Nat Biotechnol.

Gay, D., Ghinatti, G., Guerrero-Juarez, C.F., Ferrer, R.A., Ferri, F., Lim, C.H., Murakami, S., Gault, N., Barroca, V., Rombeau, I., et al. (2020). Phagocytosis of Wnt inhibitor SFRP4 by late wound macrophages drives chronic Wnt activity for fibrotic skin healing. Sci Adv 6, eaay3704.

Gonzalez-Silva, L., Quevedo, L., and Varela, I. (2020). Tumor Functional Heterogeneity Unraveled by scRNA-seq Technologies. Trends Cancer 6, 13–19.

Guerrero-Juarez, C.F., Dedhia, P.H., Jin, S., Ruiz-Vega, R., Ma, D., Liu, Y., Yamaga, K., Shestova, O., Gay, D.L., Yang, Z., et al. (2019). Single-cell analysis reveals fibroblast heterogeneity and myeloid-derived adipocyte progenitors in murine skin wounds. Nat Commun 10, 650.

Hafemeister, C., and Satija, R. (2019). Normalization and variance stabilization of single-cell RNA-seq data using regularized negative binomial regression. Genome Biol 20, 296.

Huang, D.W., Sherman, B.T., Tan, Q., Kir, J., Liu, D., Bryant, D., Guo, Y., Stephens, R., Baseler, M.W., Lane, H.C., et al. (2007). DAVID Bioinformatics Resources: expanded annotation database and novel algorithms to better extract biology from large gene lists. Nucleic Acids Res 35, W169–175.

Hughes, R.M., Simons, B.W., Khan, H., Miller, R., Kugler, V., Torquato, S., Theodros, D., Haffner, M.C., Lotan, T., Huang, J., et al. (2019). Asporin Restricts Mesenchymal Stromal Cell Differentiation, Alters the Tumor Microenvironment, and Drives Metastatic Progression. Cancer Res 79, 3636–3650.

Ikwegbue, P.C., Masamba, P., Mbatha, L.S., Oyinloye, B.E., and Kappo, A.P. (2019). Interplay between heat shock proteins, inflammation and cancer: a potential cancer therapeutic target. Am J Cancer Res 9, 242–249.

Ji, A.L., Rubin, A.J., Thrane, K., Jiang, S., Reynolds, D.L., Meyers, R.M., Guo, M.G., George, B.M., Mollbrink, A., Bergenstrahle, J., et al. (2020). Multimodal Analysis of Composition and Spatial Architecture in Human Squamous Cell Carcinoma. Cell 182, 497–514 e422.

Jiang, D., Correa-Gallegos, D., Christ, S., Stefanska, A., Liu, J., Ramesh, P., Rajendran, V., De Santis, M.M., Wagner, D.E., and Rinkevich, Y. (2018). Two succeeding fibroblastic lineages drive dermal development and the transition from regeneration to scarring. Nat Cell Biol 20, 422–431.

Jiang, Z.H., Peng, J., Yang, H.L., Fu, X.L., Wang, J.Z., Liu, L., Jiang, J.N., Tan, Y.F., and Ge, Z.J. (2017). Upregulation and biological function of transmembrane protein 119 in osteosarcoma. Exp Mol Med 49, e329.

Jin, S., Guerrero-Juarez, C.F., Zhang, L., Chang, I., Myung, P., Plikus, M.V., and Nie, Q. (2020). Inference and analysis of cell-cell communication using CellChat. bioRxiv, 2020.2007.2021.214387.

Jin, S., Guerrero-Juarez, C.F., Zhang, L., Chang, I., Ramos, R., Kuan, C.H., Myung, P., Plikus, M.V., and Nie, Q. (2021). Inference and analysis of cell-cell communication using CellChat. Nat Commun 12, 1088.

Jin, S., MacLean, A.L., Peng, T., and Nie, Q. (2018). scEpath: energy landscape-based inference of transition probabilities and cellular trajectories from single-cell transcriptomic data. Bioinformatics 34, 2077–2086.

Joost, S., Annusver, K., Jacob, T., Sun, X., Sivan, U., Dalessandri, T., Sequeira, I., Sandberg, R., and Kasper, M. (2019). The molecular anatomy of mouse skin during hair growth and rest. bioRxiv, 750042.

Jung, Y.S., Lee, H.Y., Kim, S.D., Park, J.S., Kim, J.K., Suh, P.G., and Bae, Y.S. (2013). Wnt5a stimulates chemotactic migration and chemokine production in human neutrophils. Exp Mol Med 45, e27.

Koh, D., Wang, H., Lee, J., Chia, K.S., Lee, H.P., and Goh, C.L. (2003). Basal cell carcinoma, squamous cell carcinoma and melanoma of the skin: analysis of the Singapore Cancer Registry data 1968-97. Br J Dermatol 148, 1161–1166.

Korsunsky, I., Millard, N., Fan, J., Slowikowski, K., Zhang, F., Wei, K., Baglaenko, Y., Brenner, M., Loh, P.R., and Raychaudhuri, S. (2019). Fast, sensitive and accurate integration of single-cell data with Harmony. Nat Methods 16, 1289–1296.

Kuleshov, M.V., Jones, M.R., Rouillard, A.D., Fernandez, N.F., Duan, Q., Wang, Z., Koplev, S., Jenkins, S.L., Jagodnik, K.M., Lachmann, A., et al. (2016). Enrichr: a comprehensive gene set enrichment analysis web server 2016 update. Nucleic Acids Res 44, W90–97.

Kuonen, F., Huskey, N.E., Shankar, G., Jaju, P., Whitson, R.J., Rieger, K.E., Atwood, S.X., Sarin, K.Y., and Oro, A.E. (2019). Loss of Primary Cilia Drives Switching from Hedgehog to Ras/MAPK Pathway in Resistant Basal Cell Carcinoma. J Invest Dermatol 139, 1439–1448.

La Manno, G., Soldatov, R., Zeisel, A., Braun, E., Hochgerner, H., Petukhov, V., Lidschreiber, K., Kastriti, M.E., Lonnerberg, P., Furlan, A., et al. (2018). RNA velocity of single cells. Nature 560, 494–498.

Lesack, K., and Naugler, C. (2012). Morphometric characteristics of basal cell carcinoma peritumoral stroma varies among basal cell carcinoma subtypes. BMC Dermatol 12, 1.

Lim, C.H., Sun, Q., Ratti, K., Lee, S.H., Zheng, Y., Takeo, M., Lee, W., Rabbani, P., Plikus, M.V., Cain, J.E., et al. (2018). Hedgehog stimulates hair follicle neogenesis by creating inductive dermis during murine skin wound healing. Nat Commun 9, 4903.

Liu, J., Gao, C., Sodicoff, J., Kozareva, V., Macosko, E.Z., and Welch, J.D. (2020). Jointly defining cell types from multiple single-cell datasets using LIGER. Nat Protoc 15, 3632–3662.

Lopez-Pajares, V., Qu, K., Zhang, J., Webster, D.E., Barajas, B.C., Siprashvili, Z., Zarnegar, B.J., Boxer, L.D., Rios, E.J., Tao, S., et al. (2015). A LncRNA-MAF:MAFB transcription factor network regulates epidermal differentiation. Dev Cell 32, 693–706.

Love, M.I., Huber, W., and Anders, S. (2014). Moderated estimation of fold change and dispersion for RNA-seq data with DESeq2. Genome Biol 15, 550.

Lynch, C.C., and Matrisian, L.M. (2002). Matrix metalloproteinases in tumor-host cell communication. Differentiation 70, 561–573.

Macosko, E.Z., Basu, A., Satija, R., Nemesh, J., Shekhar, K., Goldman, M., Tirosh, I., Bialas, A.R., Kamitaki, N., Martersteck, E.M., et al. (2015). Highly Parallel Genome-wide Expression Profiling of Individual Cells Using Nanoliter Droplets. Cell 161, 1202–1214.

Mielczarek-Lewandowska, A., Hartman, M.L., and Czyz, M. (2020). Inhibitors of HSP90 in melanoma. Apoptosis 25, 12–28.

Mudigonda, T., Levender, M.M., O’Neill, J.L., West, C.E., Pearce, D.J., and Feldman, S.R. (2013). Incidence, risk factors, and preventative management of skin cancers in organ transplant recipients: a review of single- and multicenter retrospective studies from 2006 to 2010. Dermatol Surg 39, 345–364.

Nguyen, Q.H., Pervolarakis, N., Blake, K., Ma, D., Davis, R.T., James, N., Phung, A.T., Willey, E., Kumar, R., Jabart, E., et al. (2018). Profiling human breast epithelial cells using single cell RNA sequencing identifies cell diversity. Nat Commun 9, 2028.

Nugent, Z., Demers, A.A., Wiseman, M.C., Mihalcioiu, C., and Kliewer, E.V. (2005). Risk of second primary cancer and death following a diagnosis of nonmelanoma skin cancer. Cancer Epidemiol Biomarkers Prev 14, 2584–2590.

Omland, S.H., Wettergren, E.E., Mollerup, S., Asplund, M., Mourier, T., Hansen, A.J., and Gniadecki, R. (2017). Cancer associated fibroblasts (CAFs) are activated in cutaneous basal cell carcinoma and in the peritumoural skin. BMC Cancer 17, 675.

Orozco, C.A., Martinez-Bosch, N., Guerrero, P.E., Vinaixa, J., Dalotto-Moreno, T., Iglesias, M., Moreno, M., Djurec, M., Poirier, F., Gabius, H.J., et al. (2018). Targeting galectin-1 inhibits pancreatic cancer progression by modulating tumor-stroma crosstalk. Proc Natl Acad Sci U S A 115, E3769–E3778.

Owen, K.L., Brockwell, N.K., and Parker, B.S. (2019). JAK-STAT Signaling: A Double-Edged Sword of Immune Regulation and Cancer Progression. Cancers (Basel) 11.

Pashirzad, M., Shafiee, M., Rahmani, F., Behnam-Rassouli, R., Hoseinkhani, F., Ryzhikov, M., Moradi Binabaj, M., Parizadeh, M.R., Avan, A., and Hassanian, S.M. (2017). Role of Wnt5a in the Pathogenesis of Inflammatory Diseases. J Cell Physiol 232, 1611–1616.

Patel, A.P., Tirosh, I., Trombetta, J.J., Shalek, A.K., Gillespie, S.M., Wakimoto, H., Cahill, D.P., Nahed, B.V., Curry, W.T., Martuza, R.L., et al. (2014). Single-cell RNA-seq highlights intratumoral heterogeneity in primary glioblastoma. Science 344, 1396–1401.

Peterson, S.C., Eberl, M., Vagnozzi, A.N., Belkadi, A., Veniaminova, N.A., Verhaegen, M.E., Bichakjian, C.K., Ward, N.L., Dlugosz, A.A., and Wong, S.Y. (2015). Basal cell carcinoma preferentially arises from stem cells within hair follicle and mechanosensory niches. Cell Stem Cell 16, 400–412.

Philippeos, C., Telerman, S.B., Oules, B., Pisco, A.O., Shaw, T.J., Elgueta, R., Lombardi, G., Driskell, R.R., Soldin, M., Lynch, M.D., et al. (2018). Spatial and Single-Cell Transcriptional Profiling Identifies Functionally Distinct Human Dermal Fibroblast Subpopulations. J Invest Dermatol 138, 811–825.

Puram, S.V., Tirosh, I., Parikh, A.S., Patel, A.P., Yizhak, K., Gillespie, S., Rodman, C., Luo, C.L., Mroz, E.A., Emerick, K.S., et al. (2017). Single-Cell Transcriptomic Analysis of Primary and Metastatic Tumor Ecosystems in Head and Neck Cancer. Cell 171, 1611–1624 e1624.

Qiu, X., Mao, Q., Tang, Y., Wang, L., Chawla, R., Pliner, H.A., and Trapnell, C. (2017). Reversed graph embedding resolves complex single-cell trajectories. Nat Methods 14, 979–982.

Rahmani, W., Abbasi, S., Hagner, A., Raharjo, E., Kumar, R., Hotta, A., Magness, S., Metzger, D., and Biernaskie, J. (2014). Hair follicle dermal stem cells regenerate the dermal sheath, repopulate the dermal papilla, and modulate hair type. Dev Cell 31, 543–558.

Rees, J.R., Zens, M.S., Celaya, M.O., Riddle, B.L., Karagas, M.R., and Peacock, J.L. (2015). Survival after squamous cell and basal cell carcinoma of the skin: A retrospective cohort analysis. Int J Cancer 137, 878–884.

Rigel, D.S., Friedman, R.J., and Kopf, A.W. (1996). Lifetime risk for development of skin cancer in the U.S. population: current estimate is now 1 in 5. J Am Acad Dermatol 35, 1012–1013.

Rinkevich, Y., Walmsley, G.G., Hu, M.S., Maan, Z.N., Newman, A.M., Drukker, M., Januszyk, M., Krampitz, G.W., Gurtner, G.C., Lorenz, H.P., et al. (2015). Skin fibrosis. Identification and isolation of a dermal lineage with intrinsic fibrogenic potential. Science 348, aaa2151.

Rowell, D., Gordon, L.G., Olsen, C.M., and Whiteman, D.C. (2016). A comparison of the direct medical costs for individuals with or without basal or squamous cell skin cancer: A study from Australia. SAGE Open Med 4, 2050312116646030.

Rubin, A.I., Chen, E.H., and Ratner, D. (2005). Basal-cell carcinoma. N Engl J Med 353, 2262–2269.

Rudolph, C., Schnoor, M., Eisemann, N., and Katalinic, A. (2015). Incidence trends of nonmelanoma skin cancer in Germany from 1998 to 2010. J Dtsch Dermatol Ges 13, 788–797.

Sahai, E., Astsaturov, I., Cukierman, E., DeNardo, D.G., Egeblad, M., Evans, R.M., Fearon, D., Greten, F.R., Hingorani, S.R., Hunter, T., et al. (2020). A framework for advancing our understanding of cancer-associated fibroblasts. Nat Rev Cancer 20, 174–186.

Sanchez-Danes, A., Larsimont, J.C., Liagre, M., Munoz-Couselo, E., Lapouge, G., Brisebarre, A., Dubois, C., Suppa, M., Sukumaran, V., Del Marmol, V., et al. (2018). A slow-cycling LGR5 tumour population mediates basal cell carcinoma relapse after therapy. Nature 562, 434–438.

Sasaki, K., Sugai, T., Ishida, K., Osakabe, M., Amano, H., Kimura, H., Sakuraba, M., Kashiwa, K., and Kobayashi, S. (2018). Analysis of cancer-associated fibroblasts and the epithelial-mesenchymal transition in cutaneous basal cell carcinoma, squamous cell carcinoma, and malignant melanoma. Hum Pathol 79, 1–8.

Satija, R., Farrell, J.A., Gennert, D., Schier, A.F., and Regev, A. (2015). Spatial reconstruction of single-cell gene expression data. Nat Biotechnol 33, 495–502.

Schindelin, J., Arganda-Carreras, I., Frise, E., Kaynig, V., Longair, M., Pietzsch, T., Preibisch, S., Rueden, C., Saalfeld, S., Schmid, B., et al. (2012). Fiji: an open-source platform for biological-image analysis. Nat Methods 9, 676–682.

Sekulic, A., Mangold, A.R., Northfelt, D.W., and LoRusso, P.M. (2013). Advanced basal cell carcinoma of the skin: targeting the hedgehog pathway. Curr Opin Oncol 25, 218–223.

Sekulic, A., Migden, M.R., Oro, A.E., Dirix, L., Lewis, K.D., Hainsworth, J.D., Solomon, J.A., Yoo, S., Arron, S.T., Friedlander, P.A., et al. (2012). Efficacy and safety of vismodegib in advanced basal-cell carcinoma. N Engl J Med 366, 2171–2179.

Sharpe, H.J., Pau, G., Dijkgraaf, G.J., Basset-Seguin, N., Modrusan, Z., Januario, T., Tsui, V., Durham, A.B., Dlugosz, A.A., Haverty, P.M., et al. (2015). Genomic analysis of smoothened inhibitor resistance in basal cell carcinoma. Cancer Cell 27, 327–341.

Sng, J., Koh, D., Siong, W.C., and Choo, T.B. (2009). Skin cancer trends among Asians living in Singapore from 1968 to 2006. J Am Acad Dermatol 61, 426–432.

Sole-Boldo, L., Raddatz, G., Schutz, S., Mallm, J.P., Rippe, K., Lonsdorf, A.S., Rodriguez-Paredes, M., and Lyko, F. (2020). Single-cell transcriptomes of the human skin reveal age-related loss of fibroblast priming. Commun Biol 3, 188.

Staples, M.P., Elwood, M., Burton, R.C., Williams, J.L., Marks, R., and Giles, G.G. (2006). Non-melanoma skin cancer in Australia: the 2002 national survey and trends since 1985. Med J Aust 184, 6–10.

Stephenson, W., Donlin, L.T., Butler, A., Rozo, C., Bracken, B., Rashidfarrokhi, A., Goodman, S.M., Ivashkiv, L.B., Bykerk, V.P., Orange, D.E., et al. (2018). Single-cell RNA-seq of rheumatoid arthritis synovial tissue using low-cost microfluidic instrumentation. Nat Commun 9, 791.

Stuart, T., Butler, A., Hoffman, P., Hafemeister, C., Papalexi, E., Mauck, W.M., 3rd, Hao, Y., Stoeckius, M., Smibert, P., and Satija, R. (2019). Comprehensive Integration of Single-Cell Data. Cell 177, 1888–1902 e1821.

Subramanian, A., Tamayo, P., Mootha, V.K., Mukherjee, S., Ebert, B.L., Gillette, M.A., Paulovich, A., Pomeroy, S.L., Golub, T.R., Lander, E.S., et al. (2005). Gene set enrichment analysis: a knowledge-based approach for interpreting genome-wide expression profiles. Proc Natl Acad Sci U S A 102, 15545–15550.

Tang, J.C., Buckel, L., and Hanke, C.W. (2019). Histopathology of Basal Cell Carcinoma After Treatment With Vismogedib. J Drugs Dermatol 18, 136–138.

Techer, H., and Pasero, P. (2021). The Replication Stress Response on a Narrow Path Between Genomic Instability and Inflammation. Front Cell Dev Biol 9, 702584.

Tellechea, O., Reis, J.P., Domingues, J.C., and Baptista, A.P. (1993). Monoclonal antibody Ber EP4 distinguishes basal-cell carcinoma from squamous-cell carcinoma of the skin. Am J Dermatopathol 15, 452–455.

Thomas, P.D., Campbell, M.J., Kejariwal, A., Mi, H., Karlak, B., Daverman, R., Diemer, K., Muruganujan, A., and Narechania, A. (2003). PANTHER: a library of protein families and subfamilies indexed by function. Genome Res 13, 2129–2141.

Tirosh, I., Izar, B., Prakadan, S.M., Wadsworth, M.H., 2nd, Treacy, D., Trombetta, J.J., Rotem, A., Rodman, C., Lian, C., Murphy, G., et al. (2016a). Dissecting the multicellular ecosystem of metastatic melanoma by single-cell RNA-seq. Science 352, 189–196.

Tirosh, I., Venteicher, A.S., Hebert, C., Escalante, L.E., Patel, A.P., Yizhak, K., Fisher, J.M., Rodman, C., Mount, C., Filbin, M.G., et al. (2016b). Single-cell RNA-seq supports a developmental hierarchy in human oligodendroglioma. Nature 539, 309–313.

Tran, H.T.N., Ang, K.S., Chevrier, M., Zhang, X., Lee, N.Y.S., Goh, M., and Chen, J. (2020). A benchmark of batch-effect correction methods for single-cell RNA sequencing data. Genome Biol 21, 12.

Trapnell, C., Cacchiarelli, D., Grimsby, J., Pokharel, P., Li, S., Morse, M., Lennon, N.J., Livak, K.J., Mikkelsen, T.S., and Rinn, J.L. (2014). The dynamics and regulators of cell fate decisions are revealed by pseudotemporal ordering of single cells. Nat Biotechnol 32, 381–386.

Van de Sande, B., Flerin, C., Davie, K., De Waegeneer, M., Hulselmans, G., Aibar, S., Seurinck, R., Saelens, W., Cannoodt, R., Rouchon, Q., et al. (2020). A scalable SCENIC workflow for single-cell gene regulatory network analysis. Nat Protoc 15, 2247–2276.

Wang, S., Drummond, M.L., Guerrero-Juarez, C.F., Tarapore, E., MacLean, A.L., Stabell, A.R., Wu, S.C., Gutierrez, G., That, B.T., Benavente, C.A., et al. (2020). Single cell transcriptomics of human epidermis identifies basal stem cell transition states. Nat Commun 11, 4239.

Wang, S., Karikomi, M., MacLean, A.L., and Nie, Q. (2019). Cell lineage and communication network inference via optimization for single-cell transcriptomics. Nucleic Acids Res 47, e66.

Wehner, M.R., Cidre Serrano, W., Nosrati, A., Schoen, P.M., Chren, M.M., Boscardin, J., and Linos, E. (2018). All-cause mortality in patients with basal and squamous cell carcinoma: A systematic review and meta-analysis. J Am Acad Dermatol 78, 663–672 e663.

Welch, J.D., Kozareva, V., Ferreira, A., Vanderburg, C., Martin, C., and Macosko, E.Z. (2019). Single-Cell Multi-omic Integration Compares and Contrasts Features of Brain Cell Identity. Cell 177, 1873–1887 e1817.

White, N.M., Masui, O., Newsted, D., Scorilas, A., Romaschin, A.D., Bjarnason, G.A., Siu, K.W., and Yousef, G.M. (2014). Galectin-1 has potential prognostic significance and is implicated in clear cell renal cell carcinoma progression through the HIF/mTOR signaling axis. Br J Cancer 110, 1250–1259.

Whitson, R.J., Lee, A., Urman, N.M., Mirza, A., Yao, C.Y., Brown, A.S., Li, J.R., Shankar, G., Fry, M.A., Atwood, S.X., et al. (2018). Noncanonical hedgehog pathway activation through SRF-MKL1 promotes drug resistance in basal cell carcinomas. Nat Med 24, 271–281.

Wolock, S.L., Lopez, R., and Klein, A.M. (2019). Scrublet: Computational Identification of Cell Doublets in Single-Cell Transcriptomic Data. Cell Syst 8, 281–291 e289.

Wysong, A., Linos, E., Hernandez-Boussard, T., Arron, S.T., Gladstone, H., and Tang, J.Y. (2013). Nonmelanoma skin cancer visits and procedure patterns in a nationally representative sample: national ambulatory medical care survey 1995-2007. Dermatol Surg 39, 596–602.

Yao, C.D., Haensel, D., Gaddam, S., Patel, T., Atwood, S.X., Sarin, K.Y., Whitson, R.J., McKellar, S., Shankar, G., Aasi, S., et al. (2020). AP-1 and TGFss cooperativity drives non-canonical Hedgehog signaling in resistant basal cell carcinoma. Nat Commun 11, 5079.

Yost, K.E., Satpathy, A.T., Wells, D.K., Qi, Y., Wang, C., Kageyama, R., McNamara, K.L., Granja, J.M., Sarin, K.Y., Brown, R.A., et al. (2019). Clonal replacement of tumor-specific T cells following PD-1 blockade. Nat Med 25, 1251–1259.

Zhang, H.M., Liu, T., Liu, C.J., Song, S., Zhang, X., Liu, W., Jia, H., Xue, Y., and Guo, A.Y. (2015). AnimalTFDB 2.0: a resource for expression, prediction and functional study of animal transcription factors. Nucleic Acids Res 43, D76–81.

Zhang, L., and Nie, Q. (2021). scMC learns biological variation through the alignment of multiple single-cell genomics datasets. Genome Biol 22, 10.

Zhao, X., Ponomaryov, T., Ornell, K.J., Zhou, P., Dabral, S.K., Pak, E., Li, W., Atwood, S.X., Whitson, R.J., Chang, A.L., et al. (2015). RAS/MAPK Activation Drives Resistance to Smo Inhibition, Metastasis, and Tumor Evolution in Shh Pathway-Dependent Tumors. Cancer Res 75, 3623–3635.

Zhao, Y., Wang, C.L., Li, R.M., Hui, T.Q., Su, Y.Y., Yuan, Q., Zhou, X.D., and Ye, L. (2017). Wnt5a promotes inflammatory responses via nuclear factor kappaB (NF-kappaB) and mitogen-activated protein kinase (MAPK) pathways in human dental pulp cells. J Biol Chem 292, 4358.

