## Supplementary Figures for "Single-cell analysis of basal cell carcinoma reveals heat shock proteins promote tumor growth in response to WNT5A-mediated inflammatory signals"

**Supplementary file**

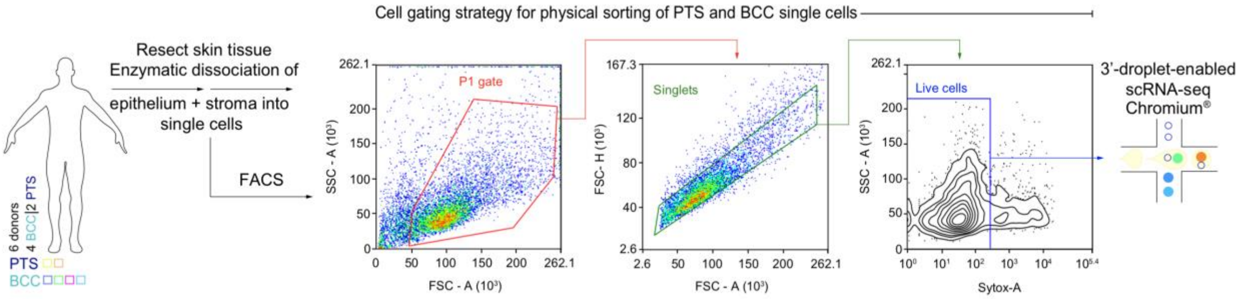


**Supplementary Figure 1. Cell sorting strategy for single cells used in 3'-droplet-enabled scRNA-sequencing.** Schematic representation of *in toto* epithelial and stromal tissue dissection, isolation, and processing from human peri-tumor skin (PTS) and basal cell carcinoma (BCC) primary clinical tumors for fluorescent activated cell sorting (FACS) and downstream 3'-droplet-enabled single-cell RNA-sequencing. Abbreviations: FACS – fluorescent activated cell sorting.

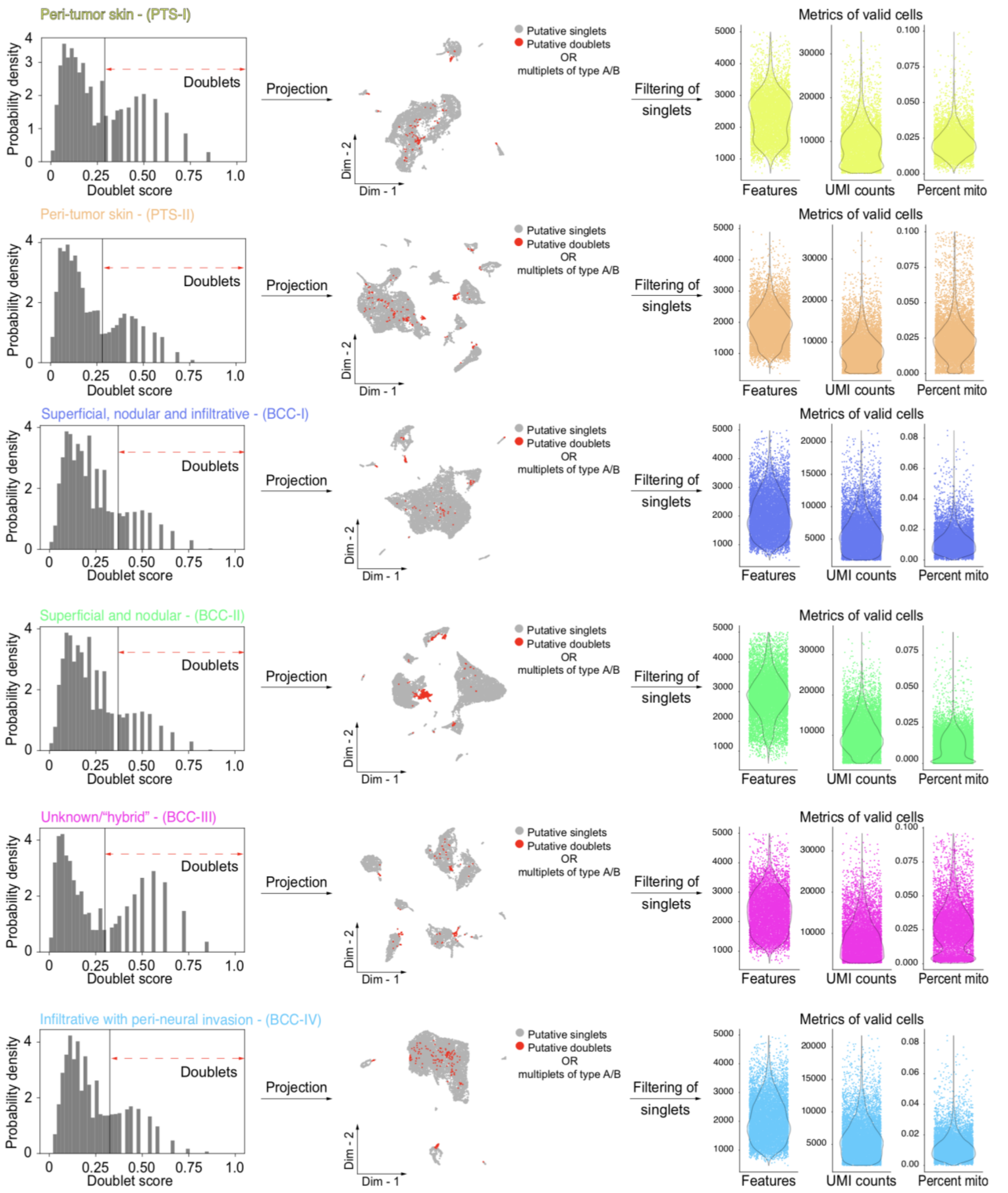


**Supplementary Figure 2. Quality control filtering of single cells.** Bimodal distribution of predicted singlet and doublet/multiplet PTS (PTS-I and PTS-II) and BCC – superficial, nodular, and infiltrative (BCC-I); superficial and nodular (BCC-II); unknown/"hybrid" (BCC-III); and infiltrative with perineural invasion (BCC-IV) cells. Straight solid line indicates user-defined doublet score threshold. Singlets were further filtered for low-quality cell pruning and removal. Putative singlets and doublets were projected onto a two-dimensional embedding and labeled accordingly. Gray indicates putative singlets; red indicates putative doublets or multiplets of type A and B. Putative singlets were processed for downstream filtering. On the right, metrics of valid cells showing distribution of features/cell, UMI counts/cell, and percentage of mitochondrial genes/cell in valid cells post-filtering are visualized as violin plots and color coded accordingly. Valid cells were used in downstream query and comparative analysis. Abbreviations: PTS – peri-tumor skin; BCC – basal cell carcinoma; FACS – fluorescent activated cell sorting.

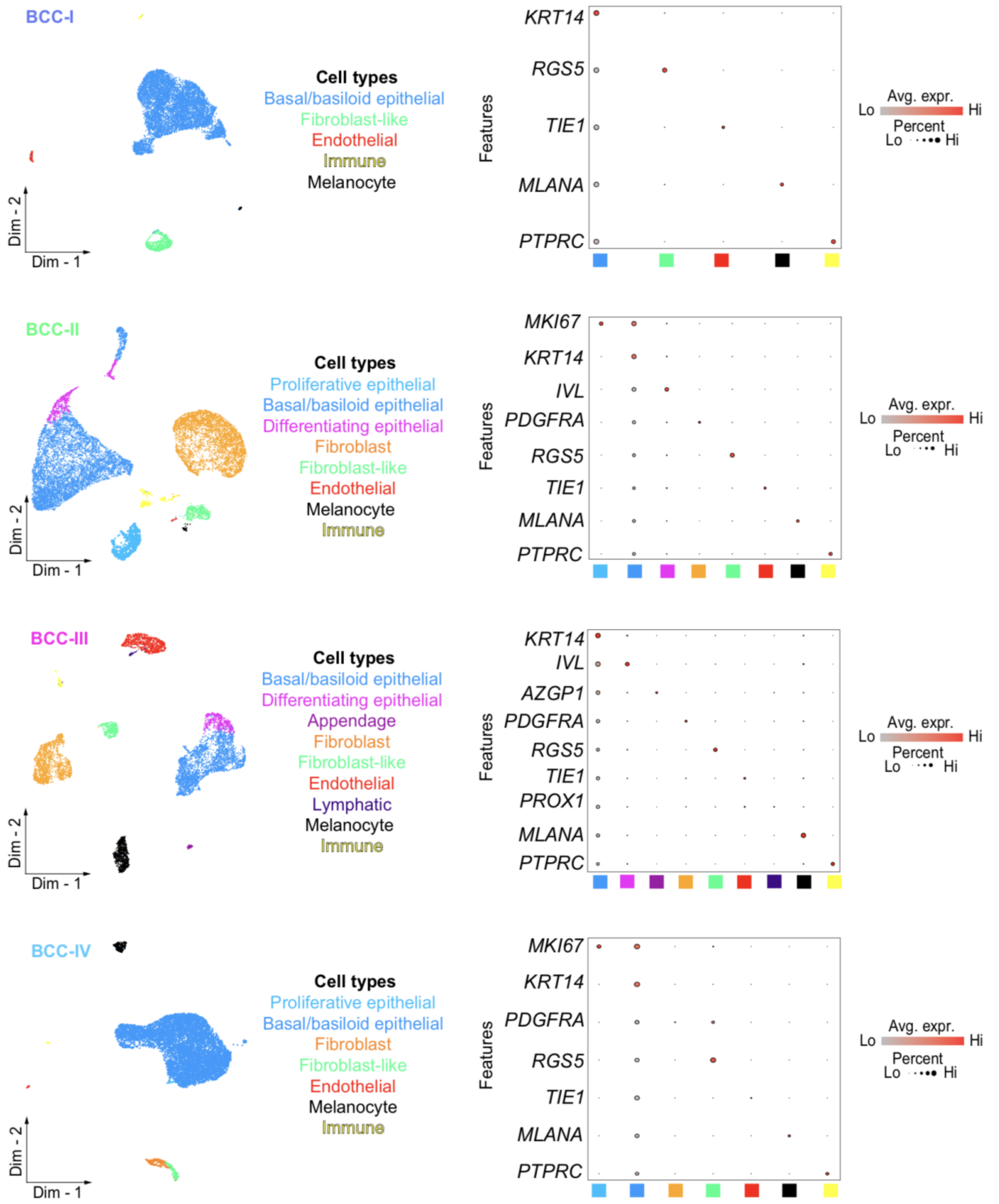


**Supplementary Figure 3. Cell type identification and characterization in basal cell carcinoma subtypes.** Two-dimensional clustering of single cells isolated from individual human basal cell carcinoma (BCC) subtypes reveals cellular heterogeneity in BCCs. IDs represent subtype and donor and are color-coded accordingly. BCC subtypes include: superficial, nodular, and infiltrative (BCC-I); superficial and nodular (BCC-II); unknown/”hybrid” (BCC-III); and infiltrative with perineural invasion (BCC-IV). Ten total distinct meta-clusters are identified at various proportions across BCC subtypes and labeled on the right. Bona fide marker for each cell type is visualized using Dotplots. Light gray indicates low average gene expression of canonical and marker genes; bright red indicates high average gene expression of canonical and marker genes. Size of circle represents the percentage of cells expressing canonical and marker genes. Abbreviations: Avg. expr. – average expression.

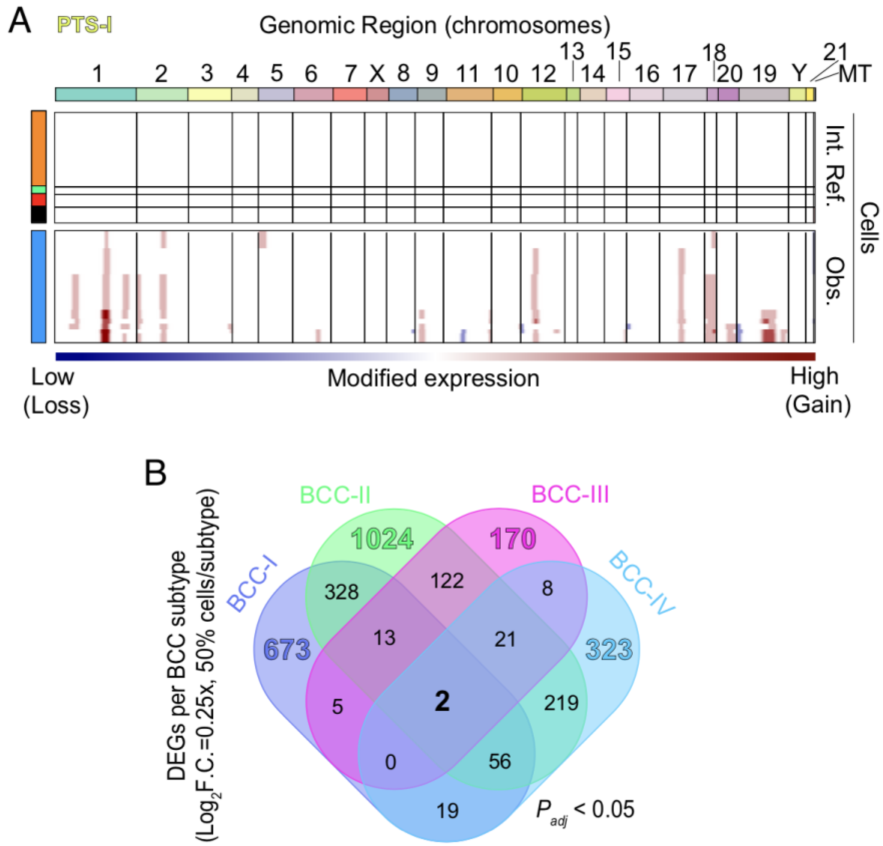


**Supplementary Figure 4. Copy number variant analysis in peri-tumor skin. A.** Copy number variant analysis of epithelial cells in peri-tumor skin (PTS-I) with InferCNV. Dark blue represents low modified expression – corresponding to genomic loss; dark red represents high modified gene expression – corresponding to genomic gain. Internal reference cells refer to non-epithelial, non-immune cells. Internal reference refers to non-epithelial control cells. Observations refer to putative malignant epithelial cells. Genomic regions (chromosomes) are labeled and color-coded. **B.** Four-way Venn diagram showing differentially expressed genes (DEGs) in BCC-I, BCC-II, BCC-III, and BCC-IV with respect to each other. DEGs per BCC subtype considered unique and significant if 50% of cells in each condition express the gene at a $Log_2$ fold change of 0.25x ($P_{adj}$ < 0.05). **I, J.** General BCC signature from this data overlaid on a two-dimensional epithelial cell embedding. Percentage of cells expressing *BCAM*, *EPCAM*, *TP63*, and *LGALS1*. Quadrupled positive cells are labeled orange and negative cells are labeled light gray. Donor-specific gene signature from this data overlaid on two-dimensional epithelial cell embedding. Percentage of cells expressing *BCAM*, *EPCAM*, *TP63*, and *LGALS1*. Quadrupled positive cells are labeled orange and negative cells are labeled light gray. Abbreviations: PTS – peri-tumor skin; BCC – basal cell carcinoma.

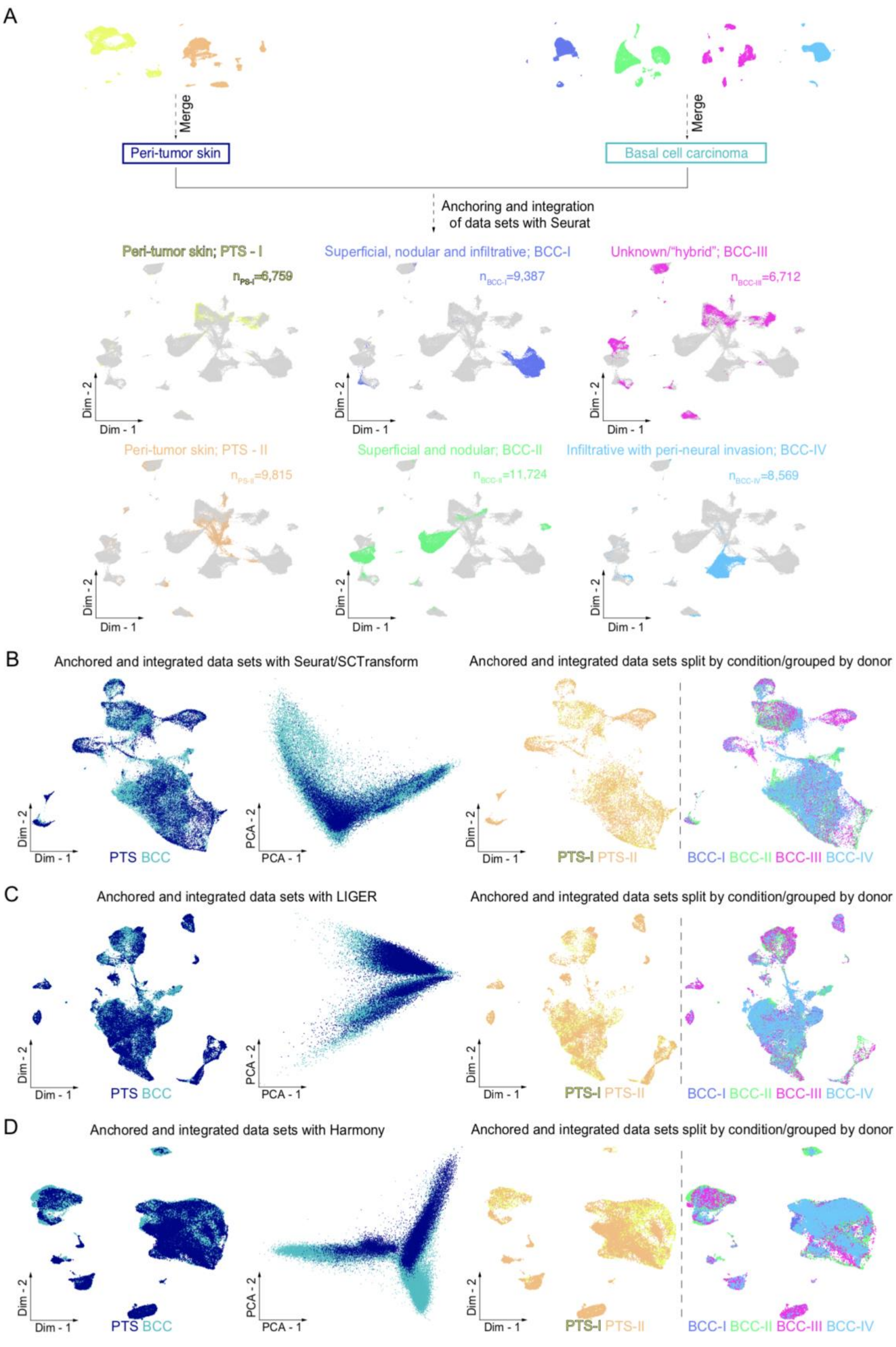

A
Merge
Merge
Peri-tumor skin
Basal cell carcinoma
Anchoring and integration of data sets with Seurat
Peri-tumor skin; PTS - I
n PS-I =6,759
Superficial, nodular and infiltrative; BCC-I
n BCC-I =9,387
Unknown/"hybrid"; BCC-III
n BCC-III =6,712
Peri-tumor skin; PTS - II
n PS-II =9,815
Superficial and nodular; BCC-II
n BCC-II =11,724
Infiltrative with peri-neural invasion; BCC-IV
n BCC-IV =8,569
Dim - 1
Dim - 2
B
Anchored and integrated data sets with Seurat/SCTransform
Anchored and integrated data sets split by condition/grouped by donor
PTS BCC
PCA - 1
PCA - 2
PTS-I PTS-II
BCC-I BCC-II BCC-III BCC-IV
C
Anchored and integrated data sets with LIGER
Anchored and integrated data sets split by condition/grouped by donor
D
Anchored and integrated data sets with Harmony
Anchored and integrated data sets split by condition/grouped by donor

**Supplementary Figure 5. Benchmarking integration of peri-tumor skin and basal cell carcinoma data sets. A-C.** Clustering of human peri-tumor skin (PTS) and basal cell carcinoma (BCC) data sets with Seurat (**A**), SCTransform (**B**), LIGER (**C**), or Harmony (**D**). Clustering was visualized with two distinct, two-dimensional embeddings. PTS and BCC data sets are color coded and split by condition and grouped by donor.

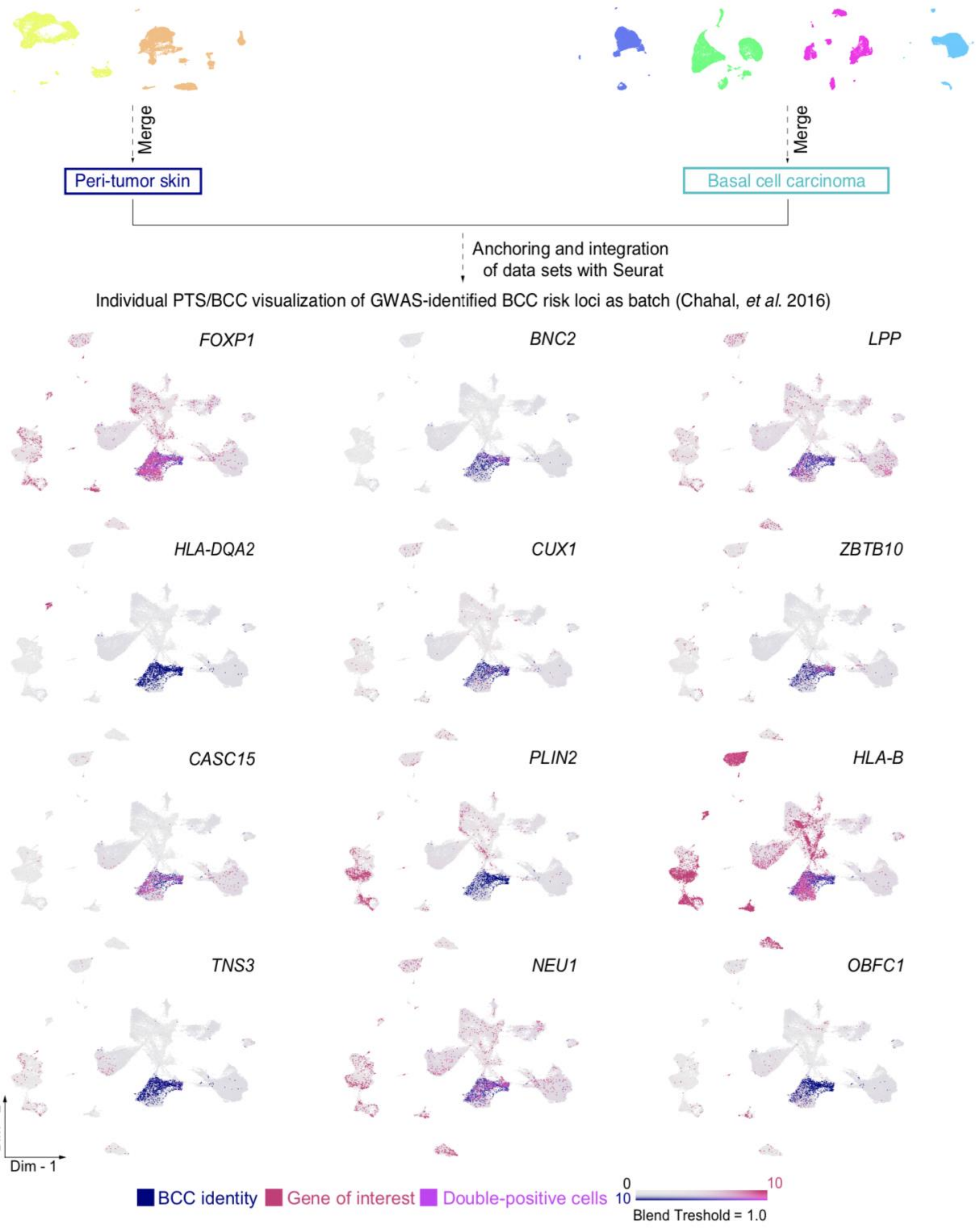


**Supplementary Figure 6. Projection of genes identified from basal cell carcinoma-risk loci GWAS analysis.** Integration of human peri-tumor skin (PTS) and basal cell carcinoma (BCC) data sets with Seurat for visualization purposes. Feature plots showing expression of basal cell carcinoma (BCC) score, gene of interest, and BCC score plus gene of interest. Gray indicates no expression; BCC-identity scored cells were colored dark blue; GWAS-identified BCC risk loci positive cells were colored red. Double-positive cells were color-coded based on a blend threshold score (blend threshold = 1.0). Abbreviations: GWAS – genome-wide associated study.

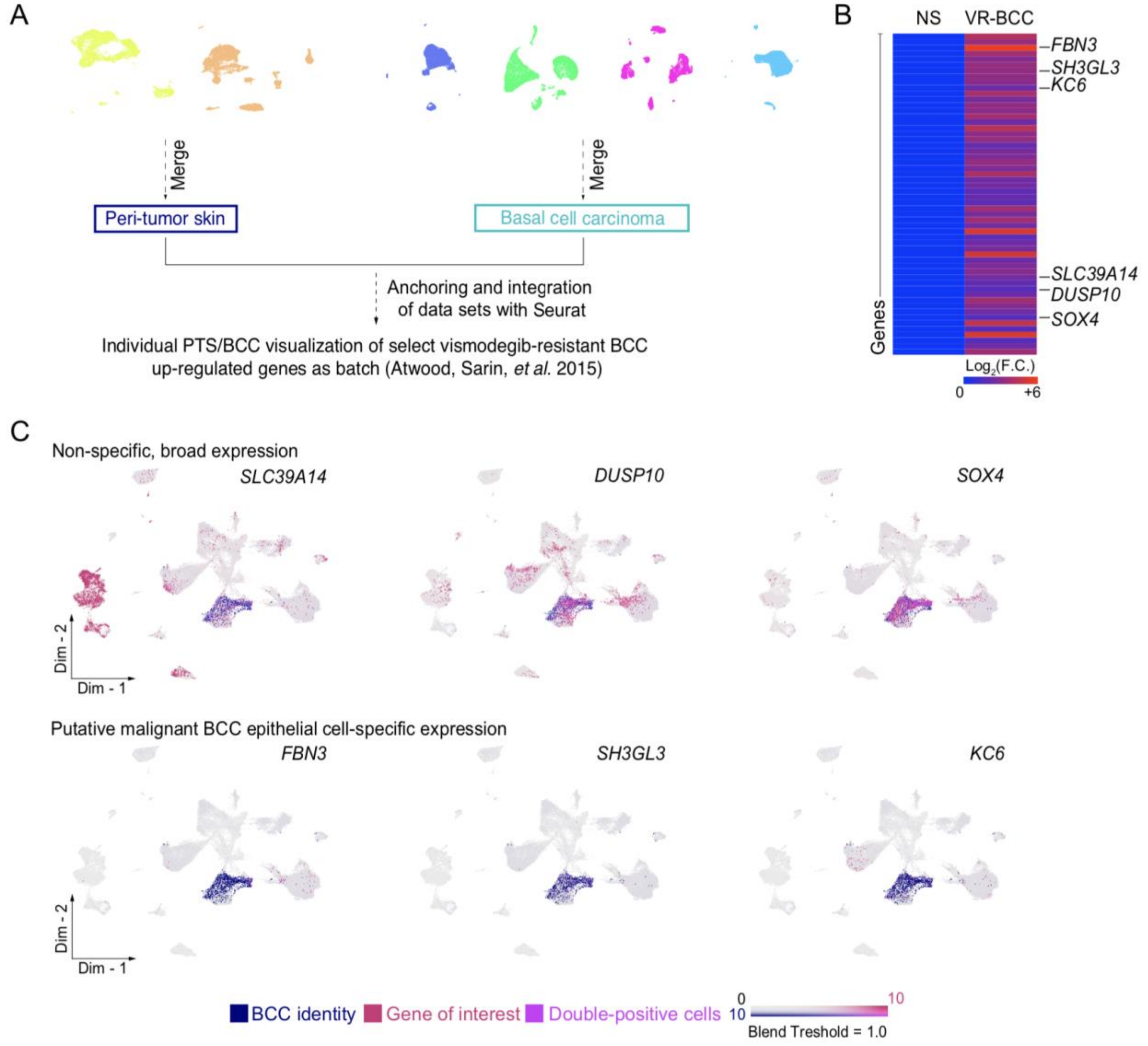


**Supplementary Figure 7. Expression of genes from bulk RNA-seq studies**. **A.** Integration of human peri-tumor skin (PTS) and basal cell carcinoma (BCC) data sets with Seurat for visualization purposes. **B.** Bulk RNA-seq analysis. Heatmap of differentially expressed genes from normal skin and vismodegib-resistant BCCs. Light blue indicates downregulation; light red indicates upregulation based on a $Log_2$ fold change. Genes of interest are on shown on the right. **C.** Feature plots showing expression of indicated marker genes categorized as "non-specific, broad expression" and "putative malignant BCC epithelial cell-specific expression". Gray indicates low expression; dark blue indicates cells which scored high for BCC genes; light red indicates cells which scored high for gene of interest; purple indicates cells which scored high for both.

Double-positive cells were color-coded based on a blend threshold score (blend threshold = 1.0). Abbreviations: NS – normal skin; VR-BCC – vismodegib-resistant basal cell carcinoma.

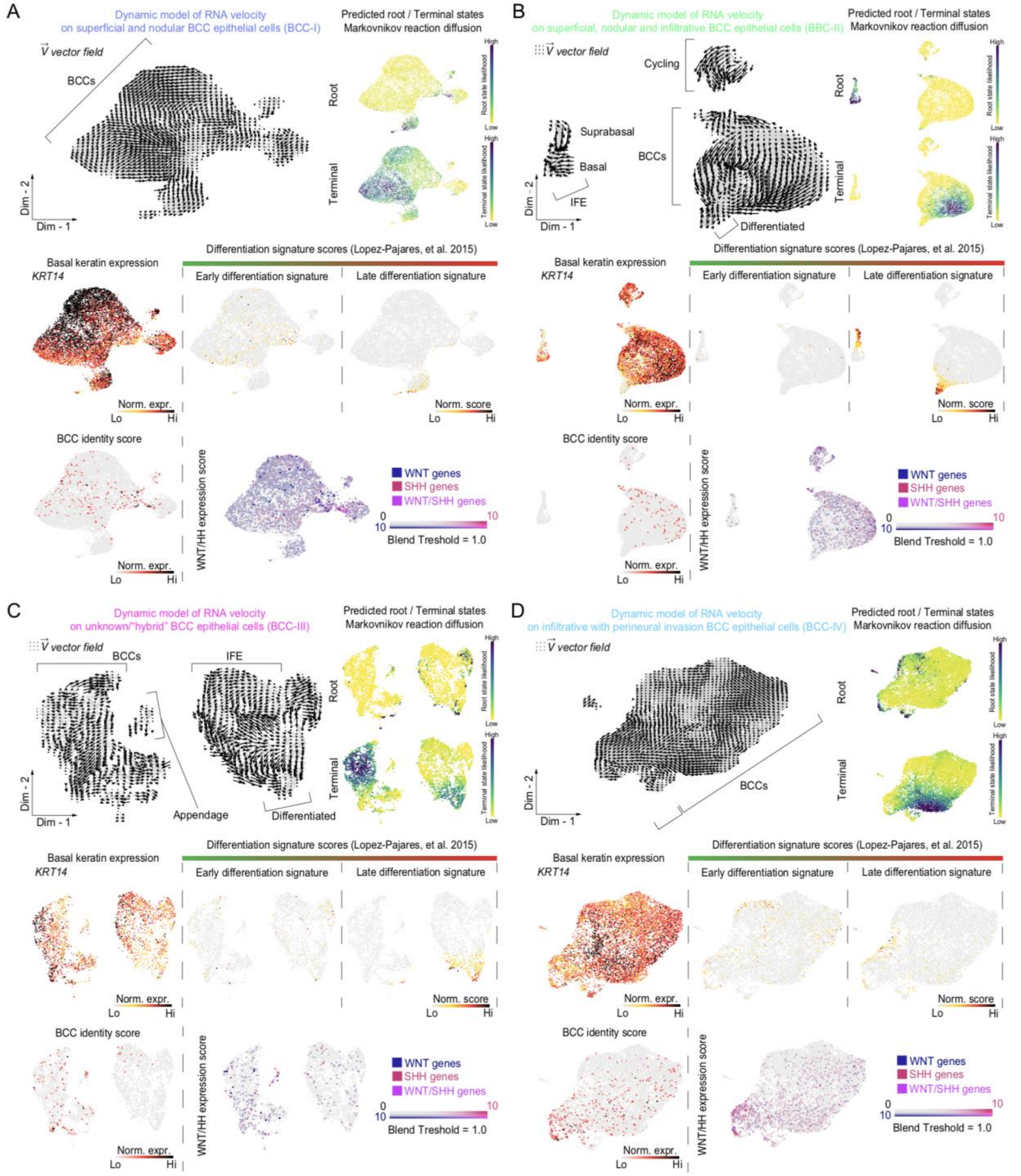


**Supplementary Figure 8. Intra-tumoral RNA dynamics analysis. A-D.** Two-dimensional sub-clustering of epithelial cells isolated from individual human basal cell carcinoma (BCC) subtypes based on basal *KRT14* expression. BCC subtypes include: (**A**) superficial, nodular, and infiltrative (BCC-I); (**B**) superficial and nodular (BCC-II); (**C**) unknown/"hybrid" (BCC-III); and (**D**) infiltrative with perineural invasion (BCC-IV). RNA velocity analysis reveals distinct intra-tumoral dynamics in BCC epithelial cells. A linear model of RNA velocity was calculated based on spliced/unspliced ratios. Resultant arrows were projected as vector field on a two-dimensional embedding. Arrows represent direction of cells' flow. Intra-tumoral predicted root and terminal states based on Markovnikov reaction diffusion are presented for each BCC subtype. Yellow indicates low probability; dark purple indicates high probability for root and terminal states. Differentiation signature scores depicting early, and late epidermal differentiation are presented for each BCC subtype. Light gray indicates low normalized gene expression based on normalized counts; black indicates high normalized gene expression based on normalized counts. BCC identity score projected on two-dimensional embedding. Light gray indicates low normalized gene expression based on normalized counts; black indicates high normalized gene expression based on normalized counts. Hedhehog (HH) and WNT-active/responsive cells were colored distinctly. Double-positive cells were color-coded based on a blend threshold score (blend threshold = 1.0). Aggregate marker gene module and blend threshold scores were Log-normalized and visualized in two-dimensional feature plots. Gray indicates no expression; dark blue indicates cells which scored high for WNT-related genes; light red indicates cells which scored high for HH-related genes; and purple indicates cells which scored high for both WNT- and HH-related genes. Abbreviations: Norm. expr. – normalized expression; Norm. score. – normalized score.

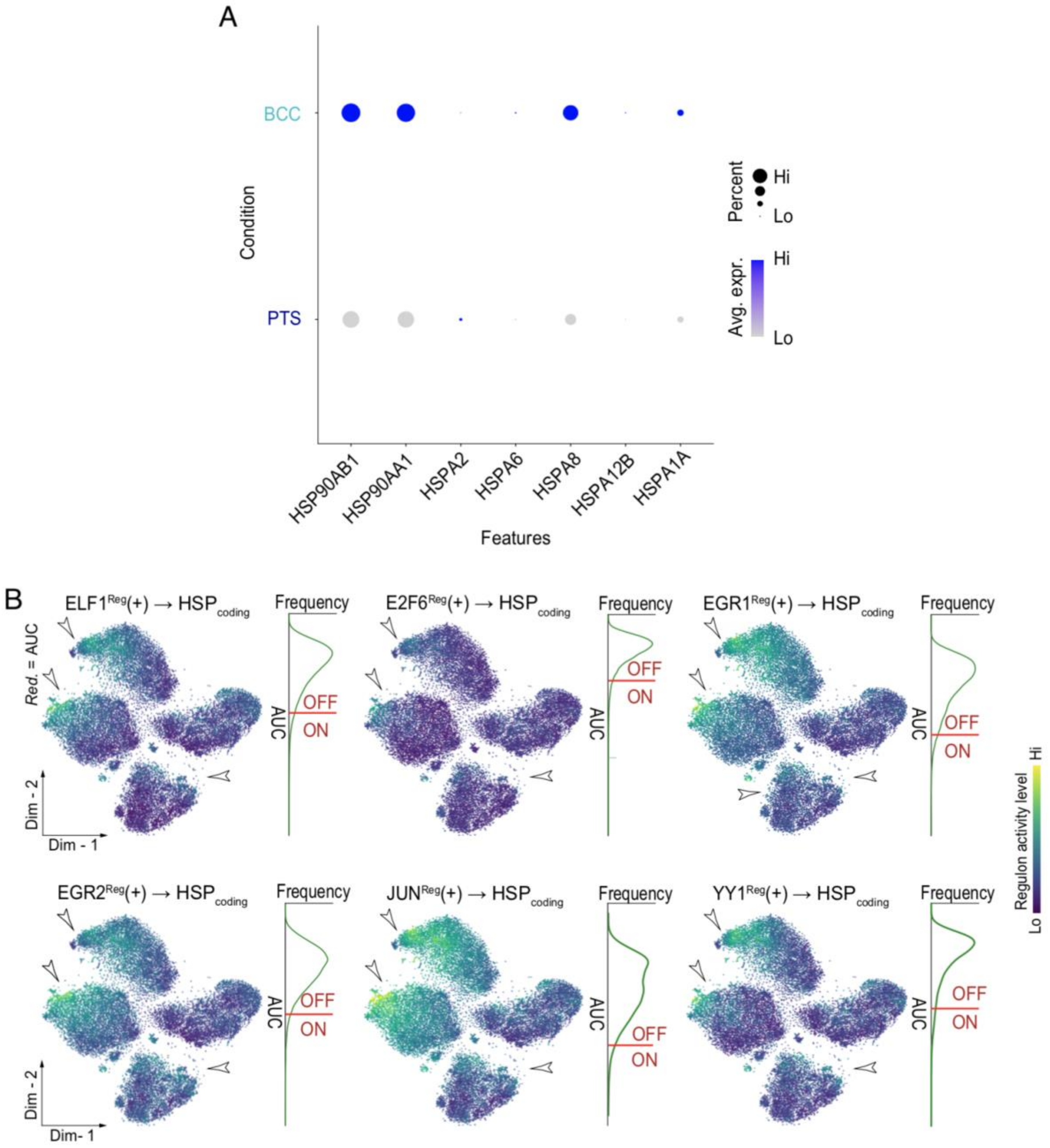


**Supplementary Figure 9. Heat shock protein-coding genes are upregulated in basal cell carcinoma. A.** Dotplot split by condition shows heat shock protein (HSP)-coding genes upregulated in human basal cell carcinoma (BCC) epithelial cells. Gray indicates low average gene expression; purple blue indicates high average gene expression. **C.** Regulon activity was used for dimensionality reduction and regulons for BCC epithelial cells were visualized in two-dimensional embedding. White arrows point at regions of high regulon activity for HSP-coding genes. Density plots represent AUC distribution per regulon selected. Dark purple indicates low regulon activity; bright yellow indicates high regulon activity. Abbreviations: PTS – peri-tumor skin; AUC – area under the curve; Avg. expr. – average expression.

**Supplementary Table S1:** Cell number quantification per individual before and after quality control (QC) filtering.

| Patient ID | Sample type | Total cells before QC | Predicted doublets | Total cells after Scrublet and QC |
|---|---|---|---|---|
| PTS-I | Peri-tumor skin | 7,164 | 170 | 6,754 |
| PTS-II | Peri-tumor skin | 10,563 | 191 | 9,777 |
| BCC-I | Primary basal cell carcinoma | 10,058 | 671 | 9,387 |
| BCC-II | Primary basal cell carcinoma | 12,511 | 787 | 11,724 |
| BCC-III | Primary basal cell carcinoma | 7,094 | 145 | 6,712 |
| BCC-IV | Primary basal cell carcinoma | 8,829 | 168 | 8,569 |

**Supplementary Table S2:** 3'-droplet-enabled single-cell RNA-sequencing metrics – mean reads per cell.

| Patient ID | Sample type | Mean reads per cell |
|---|---|---|
| PTS-I | Peri-tumor skin | 36,983 |
| PTS-II | Peri-tumor skin | 30,012 |
| BCC-I | Primary basal cell carcinoma | 48,713 |
| BCC-II | Primary basal cell carcinoma | 33,074 |
| BCC-III | Primary basal cell carcinoma | 47,337 |
| BCC-IV | Primary basal cell carcinoma | 31,405 |

**Supplementary Table S3:** 3'-droplet-enabled single-cell RNA-sequencing metrics – median genes per cell.

| Patient ID | Sample type | Median genes per cell |
|---|---|---|
| PTS-I | Peri-tumor skin | 2,382 |
| PTS-II | Peri-tumor skin | 1,902 |
| BCC-I | Primary basal cell carcinoma | 3,515 |
| BCC-II | Primary basal cell carcinoma | 2,885 |
| BCC-III | Primary basal cell carcinoma | 2,315 |
| BCC-IV | Primary basal cell carcinoma | 1,983 |
